# Phylogenetic discordance and genic innovation at the emergence of modern cephalochordates

**DOI:** 10.1101/2025.10.14.682400

**Authors:** Jaruwatana S. Lotharukpong, Christopher E. Laumer, Èlia Benito-Gutiérrez

**Author notes:** Natural History Museum, Cromwell Road, London SW7 5BD, United Kingdom (Christopher E. Laumer). Author contributions: E.B.G. and C.E.L. designed research; J.S.L., C.E.L. and E.B.G. performed research; J.S.L. and C.E.L. analysed data; J.S.L. and C.E.L. wrote the paper; E.B.G edited the paper. The authors declare no competing interests.

## Abstract

Cephalochordates (also known as amphioxus or lancelets) are a group of marine invertebrates occupying a critical phylogenetic position as the sister group to the remaining members of the chordate phylum (vertebrates and tunicates). As such, amphioxus have been key to help understanding the origin and evolution of vertebrate genomes, the evolution of their development and the diversification of their anatomy. However, out of the three genera of amphioxus currently known (*Branchiostoma, Asymmetron* and *Epigonichthys*), most studies have focused exclusively on *Branchiostoma* species. Indeed, all amphioxus genomes published to date belong to the *Branchiostoma* genus. This implies that our understanding of cephalochordate biology is limited to findings from *Branchiostoma* alone. It is consequently difficult to infer from the currently available *Branchiostoma* genomes, what aspects of their gene composition and structure are genus-specific or ancestral to cephalochordates. Here, we provide new high-quality genome assemblies and annotations from representatives of both *Asymmetron* and *Epigonichthys*. Genome-wide phylogenetic analyses reveal that, in contrast to previous mitogenomic studies, *Branchiostoma* is the sister group to *Asymmetron* and *Epigonichthys*. We further uncover discordant gene histories between these three genera, with notable asymmetry in the distribution of gene trees with minority topologies. Lastly, we find that extensive gene births and duplications preceded the origin of modern cephalochordates.

## Introduction

Extant cephalochordates (also known as amphioxus or lancelets) are represented by three genera *Asymmetron, Epigonichthys* and *Branchiostoma* and a total of around 29 species described for Branchiostoma (Poss and Boschung 1996). They occupy a critical phylogenetic position as the sister group to all other chordates (Bourlat et al. 2006; Putnam et al. 2008). Modern amphioxus species can therefore be understood as showing a mix of autapomorphic characters as well as chordate plesiomorphies no longer identifiable in other members of this clade. Since its split from the other chordates around 550 Ma (Yue et al. 2014; Delsuc et al. 2018; Subirana et al. 2020), the amphioxus has maintained relative phenotypic stasis compared to tunicates and vertebrates (Bányai et al. 2018), bearing morphological and body plan resemblance to fossil chordates from the Cambrian (Chen et al. 1995; Chen et al. 1999; Lacalli 2012). Consequently, the amphioxus has been focal to our understanding of the origin and evolution of chordates and vertebrates (Holland and Holland 2021).

Of the three genera of amphioxus, which diverged approx. 100-160 million years ago (Q.-L. Zhang et al. 2018), most studies have focused on *Branchiostoma*. For example, several *Branchiostoma* genomes have been fully assembled and annotated: *B. floridae* (Putnam et al. 2008; Simakov et al. 2020; Simakov et al. 2022), *B. lanceolatum* (Marlétaz et al. 2018; Brasó-Vives et al. 2022), *B. belcheri* (Huang et al. 2014; You et al. 2019; Huang et al. 2023) and *B. japonicum* (Huang et al. 2023). These genomes have facilitated comparative genomic studies across chordates (e.g. Putnam et al. 2008; Marlétaz et al. 2018; Brasó-Vives et al. 2022; Huang et al. 2023), as well as between metazoan phyla (e.g. Simakov et al. 2020; Simakov et al. 2022), resulting in findings such as the vertebrate-specific whole-genome duplication(s), the evolutionary relationship between the chordate subphyla, the evolution of metazoan karyotypes, and the birth and expansion of major gene families driving major vertebrate innovations. In contrast, the predominantly tropical and subtropical genera *Asymmetron* and *Epigonichthys* have received much less attention; there are only a few partial transcriptomes and mitogenome datasets (Nohara et al. 2005; Kon et al. 2007; Yue et al. 2014; Igawa et al. 2017; Subirana et al. 2020). This limited dataset may bias our understanding of the ancestral cephalochordate and, subsequently, the ancestral chordate genome towards unique features present in *Branchiostoma*.

Moreover, due to the lack of *Asymmetron* and *Epigonichthys* genomes the evolutionary relationship between the three genera remains unclear (Fig. 1A). Despite its inconsistency with morphological systematic results obtained from dozens of species (Poss and Boschung 1996), the prevailing hypothesis derived from mitogenome phylogenies places *Epigonichthys* and *Branchiostoma* as sister groups, with *Asymmetron* diverging first (Nohara et al. 2005; Kon et al. 2007; Igawa et al. 2017; Subirana et al. 2020). It has been well documented that organellar genome phylogeny may conflict with the true species phylogeny due to real discordance, since the mtDNA is inherited as a single locus. Several factors including incomplete lineage sorting (ILS) (also known as deep coalescence), introgression, gene duplication and loss, homoplastic natural selection and gene transfer can cause genes to have different evolutionary histories from that of the species (Maddison 1997; Degnan and Rosenberg 2009; Edwards 2009).

**Fig 1.**
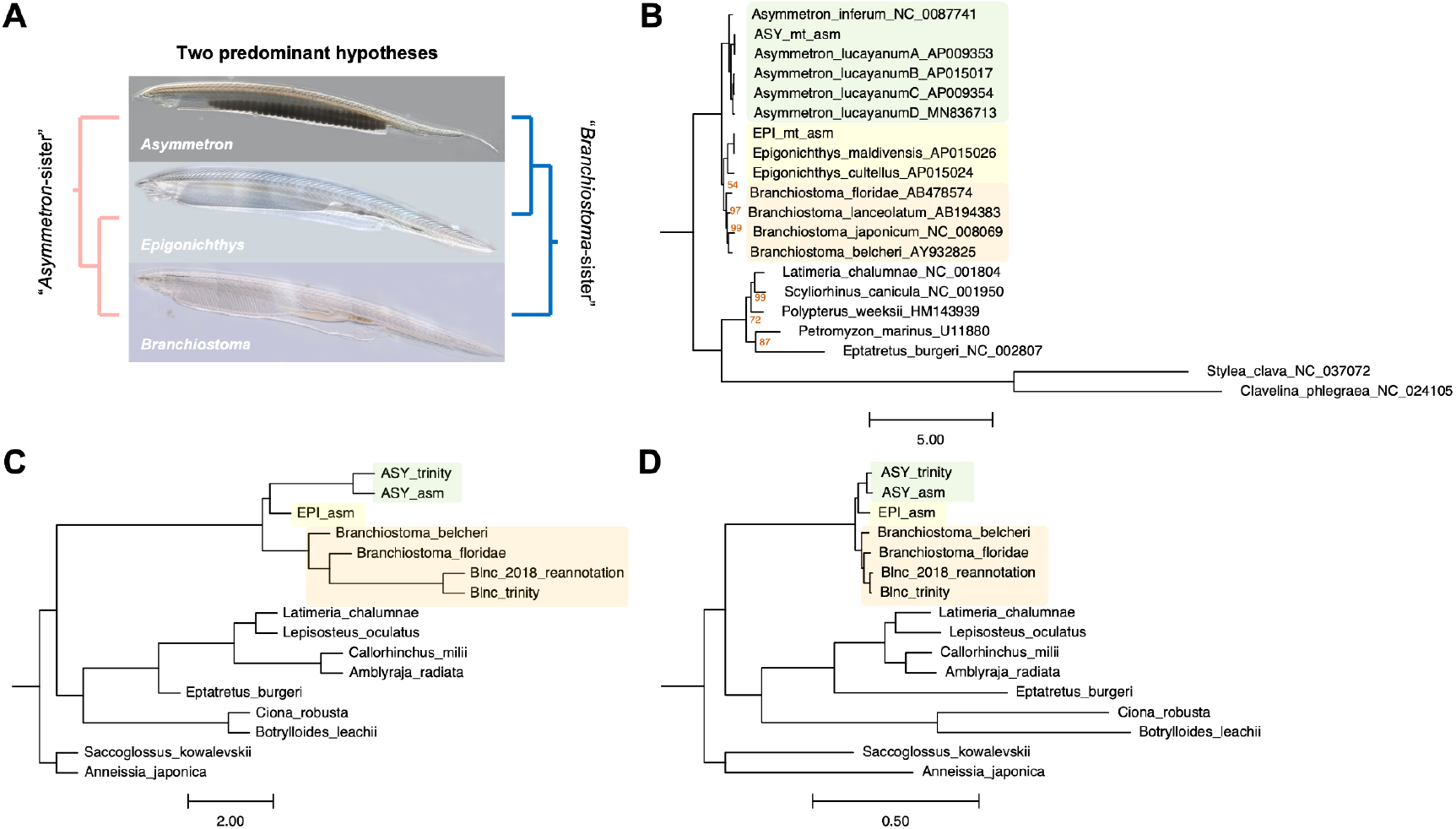
Whole-genome scale phylogenetic analysis places *Branchiostoma* as a sister group to *Asymmetron* and *Epigonichthys*, contrary to the mitogenome phylogeny. (A) Competing phylogenetic hypotheses. (B) Whole mitogenome phylogeny, including our *Epigonichthys* and *Asymmetron* specimens, confirms the monophyly of cephalochordates but fails to fully resolve relationships within the clade when applying the site-heterogeneous CAT+GTR model. (C) Quartet-based species tree analysis supports *Branchiostoma*-sister. (D) Concatenation-based maximum likelihood analysis supports *Branchiostoma*-sister. Support values are not annotated to improve visualisation, except in cases where the monophyly of the clade in question received less than maximal (100%) support.

Therefore, to advance our understanding of amphioxus biology and evolution, a whole-genome level phylogenetic analysis of these other genera is needed.

## Results

### Genome assembly and annotation

*Asymmetron* (putative *A. lucayanum* clade A) and *Epigonichthys* (putative *E. maldivensis*) isolates collected in the Maldive Islands were sequenced using Oxford Nanopore Technology (ONT) and Illumina paired-end sequencing. Prior to the genome assembly, we conducted k-mer frequency analyses on the Illumina reads using GenomeScope (Ranallo-Benavidez et al. 2020). We observed remarkably high rates of heterozygosity in both genomes (Table 1; Fig. S1). This level of heterozygosity may reflect large effective population size (Putnam et al. 2008), and can be a major hurdle for genome assembly. To overcome this and other challenges, we tested several tools and associated parameters for genome assembly, post-assembly processing, and genome annotations (Fig. S2; Tables S1, S2 & S3). Hybrid assembly approaches using MaSuRCA (Zimin et al. 2017) resulted in the best assembly statistics for both *Asymmetron* and *Epigonichthys* (Tables S1). In terms of contiguity and completeness, the *Epigonichthys* genome was the best of the two, probably because of the higher read depth coverage. The two assemblies were then purged of uncollapsed haplotype variants stemming from the high heterozygosity, polished to improve base accuracy using Illumina reads, and scaffolded using whole-individual transcriptomes (Table 1). For genome annotation, we trained the genome annotation software BRAKER2 (Brůna et al. 2021) on RNA-seq data to predict genes along the assembly scaffolds (Table 1).

**Table 1.**
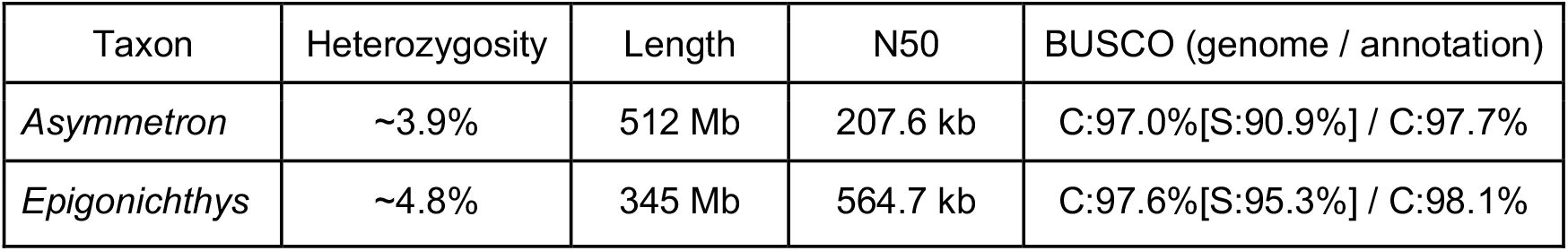
Genome statistics for the draft genomes of *Asymmetron* and *Epigonichthys*. Completeness was assessed using BUSCO metazoa_odb10.

### Revised amphioxus phylogeny

Previous mitochondrial phylogenies have suggested *Asymmetron* is the sister group to the two other genera (i.e. *Asymmetron*-sister hypothesis) (Igawa et al. 2017; Subirana et al. 2020). To repeat these results and to confirm the species assignment of the isolates we used in our assemblies, we conducted an amino-acid level phylogenetic analysis using the 13 protein-coding mitochondrial genes from the respective mitogenomes. These analyses (Figs. 1B; S3) confirm our assignment of our *Epigonichthys* isolate as *E. maldivensis* and our *Asymmetron* isolate as belonging to the Indo-Pacific “Clade A” of the *A. lucayanum* species complex.

Partitioned analysis (Fig. S3A) also recovers high, though not maximal, UF-bootstrap support values (97%) for the *Asymmetron*-sister hypothesis. However, applying mixture models under maximum likelihood (the C60 posterior mean site-frequency profile) (Wang et al. 2018) and Bayesian inference (the CAT+GTR model) erodes support for this bipartition (Figs. 1B; S3B). Moreover, a χ^2^ test of compositional heterogeneity showed all taxa but the representatives of *Asymmetron* to deviate significantly from the average amino acid frequency; average RCFV values for *Branchiostoma* and *Epigonichthys* are also 1.49 and 1.43 fold above that of *Asymmetron* (Kück and Struck 2014). We suggest, therefore, that the previously reported high support for the *Asymmetron*-sister topology in mitochondrial genome phylogenies may be an artefact of compositional non-stationarity, which is in part mitigated by the application of site-heterogeneous mixture models.

To conduct whole-genome scale phylogenetic analyses, we first collated high-quality proteomes from species that represent key groups in the deuterostome phylogeny (Table S4; Fig. S4). To fully exploit existing cephalochordate sequencing datasets, we re-assembled the transcriptome from the previously published RNA-seq dataset of *Asymmetron* (Yue et al. 2014) and re-annotated the previously published draft genome of *B. lanceolatum* (Marlétaz et al. 2018) (Table S5). We also incorporated a new genome-guided transcriptome assembly from our Mediterranean *B. lancelatum* isolate (Table S5). It should be noted that the proteome dataset contains only one isoform for each gene (Table S6). Orthology assignment, using OrthoFinder (Emms and Kelly 2019), generated 41,723 orthogroups, which contain 83.1% of 491,699 input genes (Table S7). The orthogroups were further processed to extract 1:1 orthologues for phylogenetic inference as well as to explore gene-space evolution (Fig. S5).

In contrast to the mitochondrial genome data, our whole-genome scale phylogenetic analyses (Figs. 1C & D) give maximal support for *Epigonichthys* and *Asymmetron* as sister groups to the exclusion of *Branchiostoma* (i.e. *Branchiostoma*-sister hypothesis). This is true both for a maximum-likelihood concatenated analysis of a supermatrix (length 563,157 aa with 810 partitions) (Minh et al. 2020), and for a quartet-based species tree analysis summarising 5195 gene trees (C. Zhang et al. 2018). *Branchiostoma*-sister was also supported by the paralogue-aware algorithms STAG (Emms and Kelly 2018) and ASTRAL-Pro (using 8901 gene trees) (Zhang et al. 2020), (Figs. S6A; S7).

### Gene tree discordance and potential ancient hybridisation

While the *Branchiostoma*-sister topology is consistently the best-supported, we also found high levels of gene tree discordance (Fig. 2). This discordance was observed when analysing individual gene trees inputted into the quartet-based (ASTRAL) analysis: 44.14% of gene tree quartets supported the *Branchiostoma*-sister grouping, while 31.75% and 24.15% of quartets supported the *Asymmetron*-sister and *Epigonichthys*-sister, respectively (Fig. 2A). When considering gene trees with bootstrap support over 80% (labelled as “strongly” supported), we found that ∼16% of genes strongly supported *Branchiostoma*-sister (Fig 2B). Though seemingly low, each of the alternative topologies is even less well supported in gene tree space. Analysis of the quartet-based internode certainty (Zhou et al. 2020) revealed the high discordance among gene trees, despite overall support for *Branchiostoma*-sister (Fig. S8). This contrasts with other nodes in the phylogeny such as the monophyly of cephalochordates. Furthermore, when we examined the likelihood-based phylogenetic signal in the supermatrix used in the concatenation-based analysis (computed via gene-wise log-likelihood score; see Shen et al. 2017; Shen et al. 2021), we found large proportions of gene partitions in discordance with the *Branchiostoma*-sister topology, though the weight of the distributions of phylogenetic signal is directed towards *Branchiostoma*-sister (Figs. 2C; S9).

**Fig 2.**
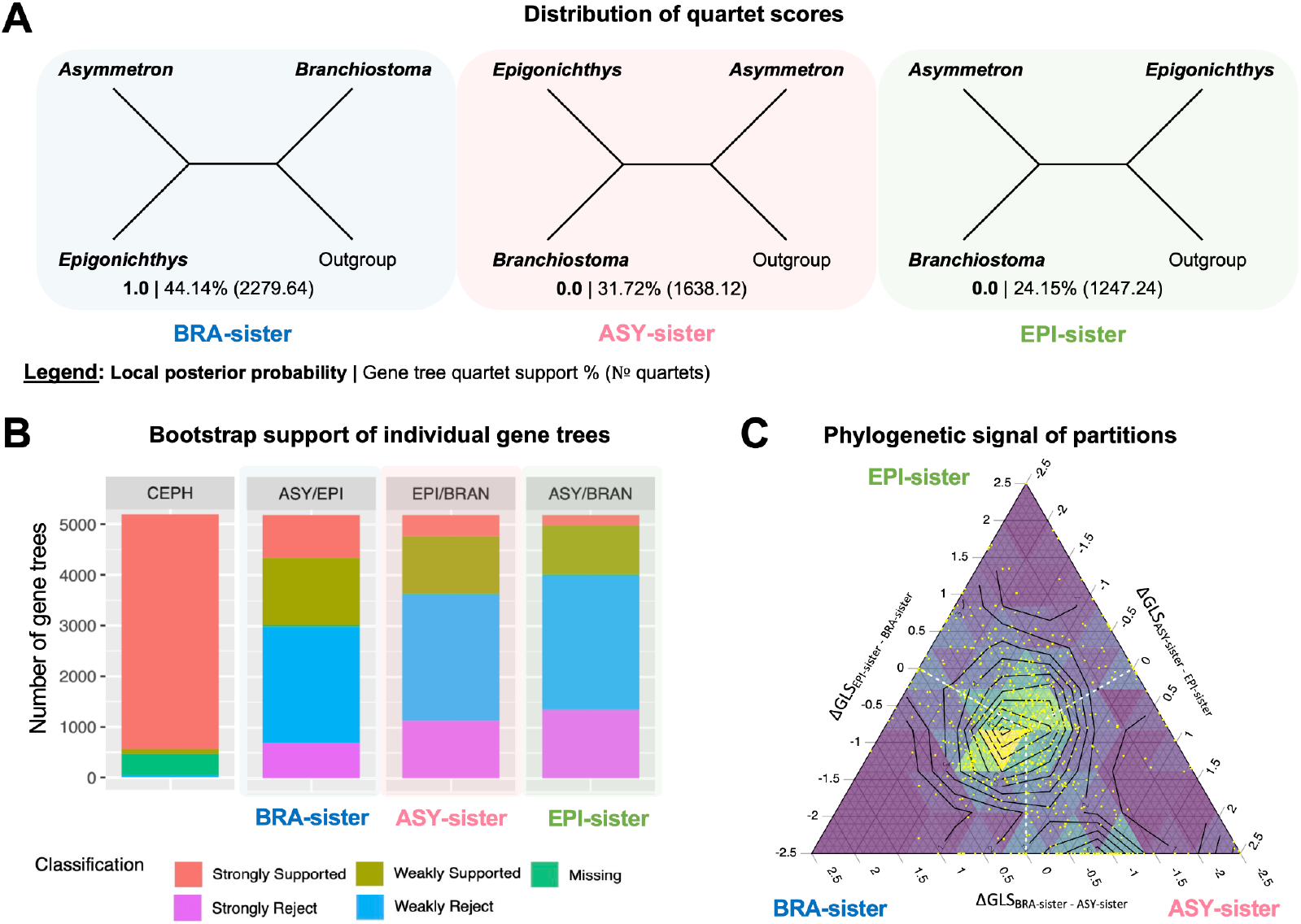
Gene tree discordance is pervasive between the three amphioxus genera, despite the support for *Branchiostoma*-sister by multigene inference. (A) The distribution of quartet scores used in the quartet-based (ASTRAL) analysis between the three possible topologies for cephalochordate genera. It should be noted that non-integer quartets can be caused by genes outside of their defined groupings. (B) Analysis of the bootstrap support of the individual gene trees that support a given grouping. (C) Analysis of the phylogenetic signal for each topology from individual partitions in the super-matrix presents a similar picture. The phylogenetic signal is computed from the difference in the average log-likelihood score supporting each topology per partition (i.e. difference in the gene-wise log-likelihood score; ΔGLS).

Gene tree discordance is not surprising given its prevalence across the tree of life due to ILS and introgression, among other factors (Degnan and Rosenberg 2009; Edwards 2009). Indeed, a third of quartets disagree with the prevailing chordate phylogeny, ((vertebrates, tunicates), cephalochordates) (Fig. S10), which reflects previous findings (Putnam et al. 2008), though the prevailing chordate phylogeny is backed with full local posterior probability support. However, what is remarkable is the degree of discordance at the base of cephalochordates. Since a rapid radiation event can lead to a genuine polytomous relationship between the genera, we conducted a polytomy analysis as implemented in PhyKit (Steenwyk et al. 2021). Polytomy at the base of the cephalochordate tree was rejected (*χ*^2^ = 308.3 with N = 5102 gene trees; p = 0.0).

Several lines of evidence (i.e. the quartet support analysis, the bootstrap support analysis and the partition analysis) reveal a disproportionate distribution of discordance between the two alternative topologies: *Asymmetron*-sister has more support than *Epigonichthys*-sister. Over 30% more gene tree quartets support *Asymmetron*-sister than *Epigonichthys*-sister (Fig. 2A). When considering the bootstrap support of individual gene trees, approximately double the number of gene trees strongly support *Asymmetron*-sister (8%) than *Epigonichthys*-sister (4%) (Fig. 2B). When comparing the phylogenetic signal of each partition, more partitions support *Asymmetron*-sister than *Epigonichthys*-sister, with the mean ΔGLS (0.285) indicating a stronger overall phylogenetic signal for *Asymmetron*-sister (Figs. 2C; S9).

### High levels of gene family birth and gene duplication in the amphioxus ancestor

We detected extensive genic innovation preceding the emergence of modern cephalochordates. Except for the origin of gnathostomes (sharks and bony fishes), cephalochordates had gene duplication levels higher than anywhere else in the deuterostome tree of life, assuming irreversibility of gene loss and duplicate retention in at least 50% of daughter taxa (Figs. 3A; S6C). Analysis using Cafe5 (which accounts for the birth-death distribution of orthogroups) indicated heightened levels of both gene family expansion and contraction in the common ancestor of modern cephalochordates compared to other groups such as vertebrates (Fig. 3A). Gnathostome origins were accompanied by a slightly higher level of gene family expansion, though the contraction was markedly lower than cephalochordates. In contrast, extensive gene family contraction preceded the origin of extant tunicates, though the level of gene family expansion is much lower than in cephalochordates. Furthermore, we extracted orthogroups exclusive to each node and further separated those represented by sequences from all daughter lineages (‘complete’) and those represented by sequences from less than 100% of daughter lineages (‘partial’). Under Dollo’s parsimony (i.e. irreversibility of orthogroup loss) (Farris 1977), we found many orthogroups to be exclusive to cephalochordates compared to other branches on the deuterostome tree (Fig. 3A), even with the conservative threshold of analysing ‘complete’ cephalochordate-exclusive orthogroups. It should be noted that RNA-seq mapping supports over 90% of gene models for at least one species within ‘complete’ cephalochordate-exclusive orthogroups, strongly suggesting these genes are not annotation artefacts, though the lack of RNA-seq mapping does not preclude evidence for the gene model (Fig. S11). The findings for gene duplication, gene family expansion/contraction and cephalochordate-exclusive gene families were robust after removing datasets with possible gene inflation (i.e. transcriptome assemblies and the re-annotated *B. lanceolatum*) (Figs. S6D; S12).

**Fig 3.**
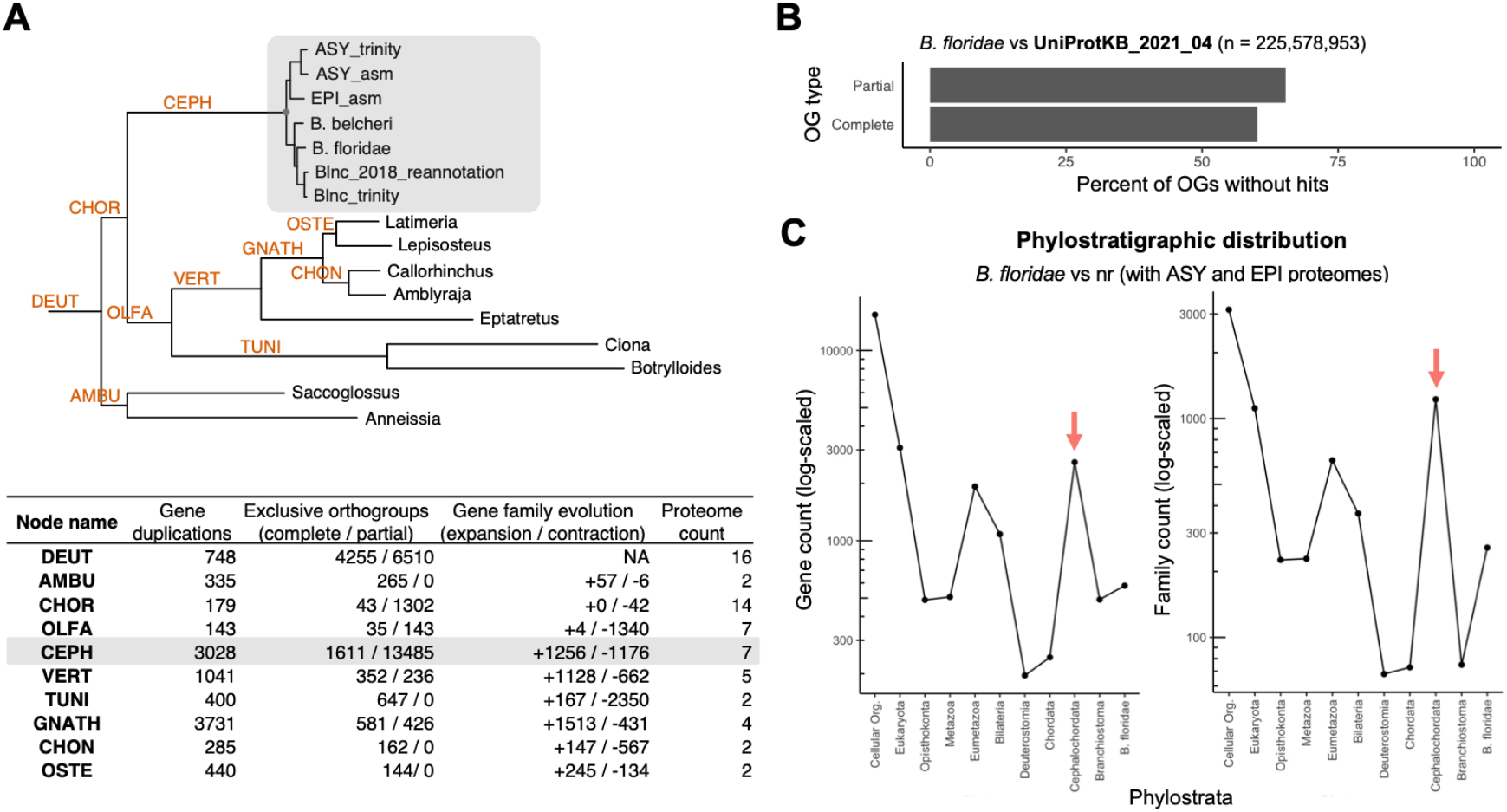
Genic innovations in the ancestor of modern cephalochordates. (A) Genic innovations at each given node inferred from the phylogenetic dataset. (B) A large proportion of the cephalochordate-exclusive orthogroups have no sequence hits outside of the group when assessed against the UniProtKB database. (C) Phylostratigraphic distribution at the individual gene and inferred gene family levels.

Phylostratigraphic analysis revealed that a majority of cephalochordate-exclusive orthogroups have no hits anywhere else in UniProtKB (DIAMOND ultra-sensitive mode with threshold pident > 30%, qcov > 70%, E-value < 1e-5) when filtered for the manually annotated of *B. floridae* (Figs. 3B; S13A & B). It should be noted that with these relatively conservative BLAST parameters, a phylogenetically novel gene may still have a hit. Using GenEra (Barrera-Redondo et al. 2023), a phylostratigraphic tool that assesses the reliability of phylostratum assignment, the resulting phylostratigraphic maps further demonstrate a burst of new genes and automatically-inferred gene families at the origin of cephalochordates (Figs. 3C; S13C & D).

These new genes represent ∼10% of the *B. floridae* genome, 20% of which were associated with gene ontology terms with the top hits being “protein binding”, “calcium ion binding”, “integral component of membrane” and “signal transduction” (Fig. S14). More gene families are exclusive to cephalochordates than other major lineages in the deuterostome phylogeny, and a large share of these families have no sequence homology to any other protein sequences in the tree of life, suggesting *de novo* origin or functional novelty.

## Discussion

### Phylogenetic analysis and gene tree discordance

Our revised phylogeny using genome-wide nuclear markers places *Branchiostoma* as the sister group to a clade consisting of the understudied tropical genera *Asymmetron* and *Epigonichthys*, contrary to previous phylogenies based on the mitogenome (Nohara et al. 2005; Kon et al. 2007; Igawa et al. 2017; Subirana et al. 2020). This topology should serve to draw experimental attention to this clade, which may retain ancestral chordate features lost during the approx. 100-160 million years ago since its split with the genus *Branchiostoma* (Q.-L. Zhang et al. 2018).

This new phylogeny also alters our understanding of morphological trait evolution in this clade. For example, the previous tree implied that the dextral-gonad placement is likely the ancestral state (Igawa et al. 2017), whereas the revised phylogeny weakens this assertion. Future studies in either two tropical genera can uncover the developmental origin of this asymmetric body plan or, alternatively, its partial loss in *Branchiostoma* and other chordates.

Based on multispecies coalescent (MSC) theory, the high degree of discordance at the origin of modern cephalochordates suggests a large effective ancestral population size or a rapid speciation event (Maddison 1997). A large effective ancestral population size is supported by the high heterozygosity rate (Putnam et al. 2008; Bi et al. 2020). A rapid speciation event has been previously reported (Igawa et al. 2017), although this is inconsistent with a formal polytomy test on our dataset (Steenwyk et al. 2021). However, under deep coalescence alone, the minority gene tree topologies are expected to have equal frequency (Degnan and Salter 2005). In contrast, we saw disproportionate phylogenetic signal favouring *Asymmetron*-sister compared to *Epigonichthys*-sister, both in raw gene tree frequency and in the distribution of resampling support within these gene trees. This has several potential explanations - the simplest, perhaps, being gene tree estimation error, for instance driven by factors such as compositional heterogeneity, as we saw evidence for in the mitogenome phylogeny. However, a biological process can also explain this asymmetry: past hybridisation events between the ancestors of modern *Epigonichthys* and *Branchiostoma* (Solís-Lemus and Ané 2016; Sayyari et al. 2018). The plausibility of this ancestral introgression hypothesis is strengthened by experimental hybridisation between different amphioxus species (Huang et al. 2023) and genera (Holland et al. 2015).

### Genic innovations at the origin of all extant amphioxus genera

Amphioxus protein sequences are slow evolving (Yue et al. 2014) and *Branchiostoma* genomes share deeply conserved macrosynteny with vertebrates (Putnam et al. 2008; Simakov et al. 2020) and even more distantly related metazoan lineages (Simakov et al. 2022). Given these observations, and the deep similarities in developmental patterning between amphioxus and other chordates (e.g. Holland et al. 1992; Andrews et al. 2021; Benito-Gutiérrez et al. 2021), to date cephalochordate genomes at large have been seen as evolutionarily conservative and therefore good proxies for the ancestral chordate genome (e.g. Marlétaz et al. 2018).

With automated bioinformatics tools, we observed a high frequency of gene duplications inferred to have been present in the last common cephalochordate ancestor, and a high frequency of gene models that cannot be placed in orthology groups outside of cephalochordates. We found extensive levels of gene duplications and gene family births, expansions and contractions preceding the origin of all modern cephalochordates. Indeed, the extent of gene duplications is only matched by branches that are thought to be preceded by whole-genome duplication events, such as the origin of vertebrates and jawed vertebrates (Ohno, 1970; Putnam *et al*., 2008; Simakov *et al*., 2020), although there is no evidence for such duplications within the cephalochordate lineage (Simakov et al. 2020). Our evidence for genome-wide gene family expansion aligns with previous studies demonstrating the expansion of developmental regulatory gene families in amphioxus, including Hairy (Minguillón et al. 2003; Jiménez-Delgado et al. 2006), *Evx* (Minguillón et al., 2002), SoxB (Meulemans and Bronner-Fraser 2007) and MyoD-Related Myogenic Regulatory Factors (Aase-Remedios et al. 2020), among others. Furthermore, a recent analysis in Branchiostoma species also indicated a prevalence of tandem gene duplications (Brasó-Vives et al. 2022). Our results suggest that this pattern was already established in the most recent common ancestor of extant cephalochordates. Furthermore, our phylostratigraphic analyses highlight a relatively large number of novel gene families unique to cephalochordates. Evidence for these cephalochordate-specific gene families has been previously observed, though not systematically examined (Guijarro-Clarke et al. 2020).

Our results further suggest that the rates of gene duplications and genic innovations are decoupled from the sequence substitution rate in cephalochordates. This contrasts with tunicates, the other group of invertebrate chordates, which display an elevated substitution rate but relatively low rates of gene duplications, gene family expansions, and gene births, as previously reported (Dehal et al. 2002; Delsuc et al. 2006; Tsagkogeorga et al. 2010; Tsagkogeorga et al. 2012; Voskoboynik et al. 2013; Guijarro-Clarke et al. 2020). Further studies are needed to determine whether cephalochordate genic innovations occurred rapidly in a short ‘pop-corn’ phase or more gradually in an extended ‘stew’ stage (Paps and Holland 2018).

However, validating these predicted gene models and understanding their biological significance will require substantial experimental work. Although the roles of these new genes and duplicates remain unclear, and how they interact with the conserved gene regulatory networks in these genomes is not yet known, the presence of numerous cephalochordate-specific gene families and duplications offers an exciting opportunity to explore how conserved gene expression and morphological stasis is maintained despite extensive genic innovation.

## Materials and Methods

### Sample collection

Specimens of Asymmetron and Epigonichthys were discovered and collected in the Maldives during the Tara Oceans expedition from 2011 to 2013. The North Sea specimen of Branchiostoma lanceolatum was obtained from coastal waters near Helgoland, Germany, through the guest scientist program at the Alfred Wegener Institute (AWI).

### Genome sequencing, assembly, post-processing and annotation

DNAs from flash-frozen specimens were extracted using a Qiagen Genomic Tips kit following the manufacturer’s protocol. To generate ONT reads, we used the LSK108 ligation kit to prepare libraries on unsheared DNAs, using two MinION R9.4.0 flow cells per specimen for sequencing and Guppy v4.0.14 for HAC base-calling. The Illumina paired-end reads were sequenced using a single HiSeq 4000 2x150 lane. For the Illumina reads, we used GenomeScope v2.0 (Ranallo-Benavidez et al. 2020) to estimate the genome size, abundance of repetitive elements, heterozygosity and other genome content information using a k-mer count histogram produced by Jellyfish v2.3.0 (Marçais and Kingsford 2011) (with k-mer length `-m 21` as recommended by GenomeScope). Based on these genome size estimates, the ONT sequencing depth coverages for *Asymmetron* and *Epigonichthys* are 21× and 40×, respectively. The Illumina sequencing depth coverages for *Asymmetron* and *Epigonichthys* are 126× and 164×, respectively.

For the assembly and post-processing of both *Asymmetron* and *Epigonichthys* genomes, we tested several tools and their associated parameters. The final draft genome of *Asymmetron* used in this study was assembled using hybrid assembler MaSuRCA (Zimin et al. 2017) and a custom run of Flye v2.8.2 (Kolmogorov et al. 2019). To retain all ONT reads in the assembly, MaSuRCA was run with `LHE_COVERAGE=60`. The intermediate ‘mega-reads’ output of MaSuRCA was inputted into Flye v2.8.2 through `-nano-corr`, with adjusted minimum overlap between reads (`-m 2500`) and no polishing interaction (`-i 0`). Polishing was done using POLCA (Zimin and Salzberg 2020), with default parameters. Erroneous haplotigs were purged using purge_dups (Guan et al. 2020). For this, BWA-MEM (Li 2013) was used to map the Illumina reads. The read depth cutoffs from `ngscstat` were manually curated and `get_seqs` was used without `-e` option to allow haplotypic duplicate removal from the middle of contigs.

Repeats were masked using RepeatModeler v2.0.1 (Flynn et al. 2020) and RepeatMasker v4.1.2 (www.repeatmasker.org) from the TETool v1.3 docker container. The option `-LTRStruct` was included to enable the LTR structural finder, and soft-masking was performed with `-xsmall`. The contigs were scaffolded using P_RNA_scaffolder on RNA-seq reads trimmed using TRIMMOMATIC (Bolger et al. 2014). TRIMMOMATIC was run with `ILLUMINACLIP:TruSeq2-PE.fa:4:30:10 SLIDINGWINDOW:5:15`. These trimmed reads were first mapped onto the hard-masked genome using hisat2 (Kim et al. 2019: 2), which was run with `-k 3 -pen-noncansplice 1000000`. P_RNA_scaffolder was run on the output of hisat2 and the soft-masked assembly. The `P_RNA_scaffolder.sh` code was edited to include `-masked=soft` parameter for the pairwise sequence aligner BLAT (Kent, 2002) to avoid scaffolding using repetitive element transcripts. The final draft genome of *Epigonichthys* was generated in the same manner except during the assembly step, where the final output of MaSuRCA (with `LHE_COVERAGE=60`) was used. BRAKER ET (Brůna et al. 2021) was used with `--softmasking` for both *Asymmetron* and *Epigonichthys* annotations, such that the gene predictions were aided using trimmed RNA-seq as evidence for genes. The final assemblies and annotations are labelled in the analysis as “ASY_asm” and “EPI_asm”. The same annotation method was used for the re-annotation of *B. lanceolatum* genome (from Marlétaz et al. 2018), labelled in the analysis as “Blnc_2018_reannotation”.

In addition, we assembled the genome of the North Sea specimen of *B. lanceolatum*, however, this genome assembly was not used in downstream orthology and phylogenetic analyses due to its lower quality genome annotation results (BUSCO metazoa_odb10 C < 85%), despite the genome’s high contiguity and completeness (length 482 Mb, N50 7 Mb, BUSCO eukaryota_odb10 C:99.3%[S:97.3%], BUSCO metazoa_odb10 C:96.0%[S:95.2%]). In brief, this genome was first assembled from ONT reads using canu v2.1.1 (Koren et al. 2017), with parameters for keeping haplotypes (`corOutCoverage = 200 “batOptions = -dg 3 –db 3 -dr 1 -ca 500 -cp 50”`). Post-assembly polishing was done with Racon v1.4.3 (Vaser et al. 2017), using default parameters. Haplotig-purging was done through purge_dups with Canu trimmed reads, manually set cutoffs, minimap2 parameter `-xasm20` and without `-e` for `get_seqs`. Other post-processing steps are the same as done for *Asymmetron* and *Epigonichthys*.

Genome assemblies and raw data are available in INSDC under BioProject XXXXXX.

### Transcriptome sequencing and assembly

Total RNA was isolated from flash-frozen specimens, 3 adult individuals from each genera, following homogenisation in TRIzol reagent. Phase separation was achieved with Heavy Phase Lock Tubes following the manufacturers instructions (5prime). RNA was precipitated with isopropanol, washed in cold 75% ethanol and resuspended in RNAse-free water. RNA was further purified after precipitation with RNA cleanup columns from Qiagen before library preparation. 100bp paired end libraries were prepared and sequenced by Genoscope using the Illumina TruSeq RNA kit and a HigSeq2000 sequencer,, resulting in ∼400 million reads per pair for *Asymmetron* and ∼360 Million reads per pair for *Epigonichthys*, which were used for genome annotation.

The transcriptomes of *A. lucayanum* (SRX437621) and *B. lanceolatum* were assembled using Trinity (Grabherr et al. 2011; Grabherr et al. 2011; Haas et al. 2013), with the genome-free mode and genome-guided mode, respectively. These are labelled “ASY_trinity” and “Blnc_trinity”. The genome-guided assembly was enabled through `--genome_guided_bam` and the genome assembly of the North Sea specimen of *B. lanceolatum* guided the Blnc_trinity assembly. RNA-seq reads were mapped using hisat2 for the Trinity assembly, for which the maximum intron size was set at 10 kb (`--genome_guided_max_intron 10000`).

### Mitochondrial genome assembly and phylogeny

Mitochondrial genomes from our *Epigonichthys maldivensis* and *Asymmetron lucayanum* Clade A isolates were assembled from the Illumina reads using GetOrganelle, using COI sequences identified using BLAST (Altschul et al. 1997). Mitogenomes, including those downloaded from NCBI, were annotated in MITOS2, using the RefSeq Metazoa 89 database as an annotation reference and selecting genetic codes 2 or 5 as appropriate. The annotated FASTA files were then manually stripped of any partial gene duplicates (present for nad5 in *A. inferum* and *A. lucayanum* clade D) and parsed into multi-sequence fastas using a custom bash script.

Sequences were aligned with the MAFFT L-INS-i algorithm and concatenated without trimming using catsequences (Creevey and Weeks 2021). Maximum likelihood inference using the PMSF mixture model approximation was carried out in IQ-Tree2, setting “`-m MtZOA+C60+F+G -B 1000 -bnni`”. Partitioned ML inference proceeded using an initial partition-finding test with the “`-m MF+MERGE`” and `“-rcluster 10`” options, followed by full ML inference using the best-found partition scheme and “`-B 1000 -bnni`” as bootstrapping options. Phylobayes CAT+GTR inference was carried out in pb_mpi v1.8 using “`-cat -gtr -dc`” as options across four replicate chains run for at least 7936 generations; two chains were summarised together with “`bpcomp -x 2000 10`”, giving a maxdiff value of 0.0525416, consistent with convergence of the posterior distributions.

### Species tree inference

The phylogenetic dataset was constructed using our *Asymmetron* and *Epigonichthys* genome annotations, as well as the transcriptome assemblies. Publicly available proteomes were downloaded as indicated in Table S4. Longest isoforms were retained for each gene using the scripted provided by KinFin (Laetsch and Blaxter 2017), before running OrthoFinder v2.5.2 (Emms and Kelly 2019) with 64 threads to infer orthogroups and hierarchical orthogroups at node 0 (which includes all taxa). For the transcriptome assemblies, the longest isoforms and subsequently the longest ORFs were extracted using a script provided by Trinity v2.12.0 and TransDecoder v5.5.0 (https://github.com/TransDecoder/TransDecoder), respectively.

Orthogroups were analysed to infer gene duplications, gene (sub-)family births and expansion/contraction. KinFin v1.0 (Laetsch and Blaxter 2017) and Gephi v0.9.2 (Bastian et al. 2009) were used to visualise the orthogroups (Fig. S4). Meanwhile, hierarchical orthogroups at node 0 were used to construct the phylogeny. OrthoFinder’s default phylogeny was constructed using STAG (Emms and Kelly 2018) and rooted using STRIDE (Emms and Kelly 2017), while default MSA phylogeny was inferred through the `-msa` option where the MSA was constructed using MAFFT v7.480 and FastTree v2.1.10 (Price et al. 2010). Columns with more than 50% gaps were trimmed. The hierarchical orthogroups were further filtered with a custom bash script by the following criteria: each hierarchical orthogroup must include every amphioxus taxon and at least one non-amphioxus taxon.

For both the concatenation-based IQ-TREE and quartet-based ASTRAL analyses, the 8902 hierarchical orthogroups were pruned in order to obtain, for each orthogroup, only sequences inferred to be orthologues. We first constructed MSAs using einsi and gene trees were inferred by running IQ-TREE v2.1.2 (Minh et al. 2020) on the 8901 successfully completed MSAs with IQ-TREE parameters `--seqtype AA -keep-ident -B 1000`. ModelFinder was used to automatically select the best substitution model (Kalyaanamoorthy et al. 2017). The gene trees were pruned and monophyletically-masked using PhyloPyPruner v1.2.3 (https://gitlab.com/fethalen/phylopypruner). PhyloPyPruner parameters were set at `--min-len` `50 --trim-lb 5 --min-support 0.75 --prune MI --min-taxa 8 --min-otu-occupancy 0.1 --mask pdist`. *Anneissia* and *Saccoglossus* were selected as outgroup taxa with `--outgroup`. This resulted in 5196 pruned trees as well as a concatenated supermatrix of length 4.5 M amino acids. For the concatenation-based IQ-TREE analysis, the supermatrix was post-processed using MARE v0.1.2-rc (Misof et al. 2013), to reduce the size of the concatenated supermatrix. Successful reduction to 563K amino acids was achieved through MARE parameters `-d 30.0 -t 100`. On this reduced supermatrix, partitioned ML analysis was performed (with IQ-tree2 options `-m MFP-MERGE -alrt 1000 -B 1000 --seqtype AA`). For the quartet-based whole-genome phylogeny, we ran ASTRAL v5.7.7 (C. Zhang et al. 2018) on the 5195 pruned gene trees, using option `-t 2` for full annotation. ASTRAL-Pro v1.1.4 (Zhang et al. 2020) was also used on all 8901 gene trees to incorporate out-paralogues and in-paralogues. The resulting species trees were visualised using TreeViewer v2.0.1 (https://github.com/arklumpus/TreeViewer).

### Gene tree discordance analysis

For the quartet-based discordance analysis, QuartetScore was used to compute quartet-based internode certainty (IC) scores on the input pruned trees. Gene tree discordance was visualised through DiscoVista (Sayyari et al. 2018), with analysis type option `-m 1`. The 80% bootstrap threshold for “strongly supported” and “strongly rejected” was specified through `-t 80`. In short, bootstrap support higher than 80% is binned as “strongly” supported. Gene trees that reject the given topology are binned as “weakly rejected” if compatibility is recovered with the contraction of low support branches below 80% bootstrap support. The polytomy analysis was conducted using Phykit (Steenwyk et al. 2021).

For the concatenation-based IQ-TREE analysis, the likelihood-based signal (i.e. gene-wise log likelihood score) for each partition between the main and alternative topologies were examined (as implemented in Shen et al. 2017; Shen et al. 2021). IQ-TREE was run as previously stated with additional parameters, `-z` to input the main and two alternative topologies and `-wsl` to output the site log-likelihoods in TREE-PUZZLE format. The perl script GLS_parser_v1.pl (Shen et al. 2021) was used to calculate the ΔGLS (the difference in gene-wise log likelihood score between two topologies). The ternary plots were made using the Ternary package (Smith 2022) in R.

### Gene duplication and family expansion/contraction estimation

OrthoFinder was used to estimate the number of gene duplications at each node in the deuterostome phylogeny with at least 50% support at the terminal nodes. Cafe v5.0 (Mendes et al. 2020) was used to infer the gene (sub-)family expansion/contraction. To generate the input for the Cafe5 analysis, the concatenation-based IQ-TREE phylogeny was rooted, ultrametricised (correlated model) and dated (deuterostome between 530 Ma and 636 Ma) using the ape package in R (Paradis and Schliep 2019). For Cafe5 analysis with default parameters (lambda with no among family rate variation), 100 orthogroups were removed due to large family size differentials. We also tested the consequences of allowing family rate variation with three discrete gama rate categories (`-k 3`), as well as removing taxa with possibly inflated numbers of genes (e.g. transcriptome assemblies), which did not require any orthogroup removal (Fig. S12). These results are congruent.

### Phylostratigraphy analysis

For the orthogroup phylostratigraphy, the number of new orthogroups that emerged at each node in the phylogeny was inferred using the Dollo Parsimony algorithm embedded in KinFin (Laetsch and Blaxter 2017). Orthogroups that appear at the cephalochordate node were segregated based on 100% presence in daughter lineages (‘complete’) and <100% presence (‘partial’). Gene models belonging to these orthogroups were assessed for RNA-seq mapping evidence using the BRAKER script `selectsupportedsubsets.py`. Genes within the anySupport category were selected. The accelerated-BLAST algorithm, DIAMOND (Buchfink et al. 2021), was used to detect orthogroups with hits outside the cephalochordate group. Filtering for *B. floridae* (BFLOR) sequences was done to ensure the inclusion of only high-quality gene models aided by manual curation. The DIAMOND search was conducted with the ultra-sensitive mode and the thresholds pident > 30%, qcov > 70% and E-value < 1e-5. GenEra (Barrera-Redondo et al. 2023) was used to infer the phylostratigraphic distribution of *B. floridae* genes. Alongside the NCBI NR database (accessed on 14.04.2022), the target database included the newly assembled and annotated *Asymmetron* and *Epigonichthys*. The same analysis was performed for *Asymmetron* and *Epigonichthys* genes. Default parameters were used. Gene families were automatically inferred using the Markov cluster algorithm, MCL (Enright et al. 2002), which is embedded in GenEra. Interproscan v5.54-87.0 was used to report the Gene Ontology terms of novel cephalochordate genes.

The scripts used for the analysis are documented in https://github.com/LotharukpongJS/Cephalogenomics.

## Acknowledgement

We thank John Marioni for facilitating access to the HPC cluster at the EMBL-EBI where the analysis was performed. Kevin Kocot provided a bash script we used for mitogenome phylogenetic data curation. Josué Barrera-Redondo provided guidance on phylostratigraphy. We thank Mark Blaxter for careful reading of the manuscript and for valuable feedback and support. We also thank David E. K. Ferrier, Richard Durbin, John Welch, and members of the Benito-Gutiérrez group (Giacomo Gattoni, Michael A. Schwimmer, Daniel Keitley and Christo Christov) who supported this study through scientific discussion.

