## Supplementary Information for "Phylogenetic discordance and genic innovation at the emergence of modern cephalochordates"

### Supplementary figures and tables for *Phylogenetic discordance and genic innovation at the emergence of modern cephalochordates*


Supplementary figures

Supplementary tables

Bibliography

### Supplementary figures

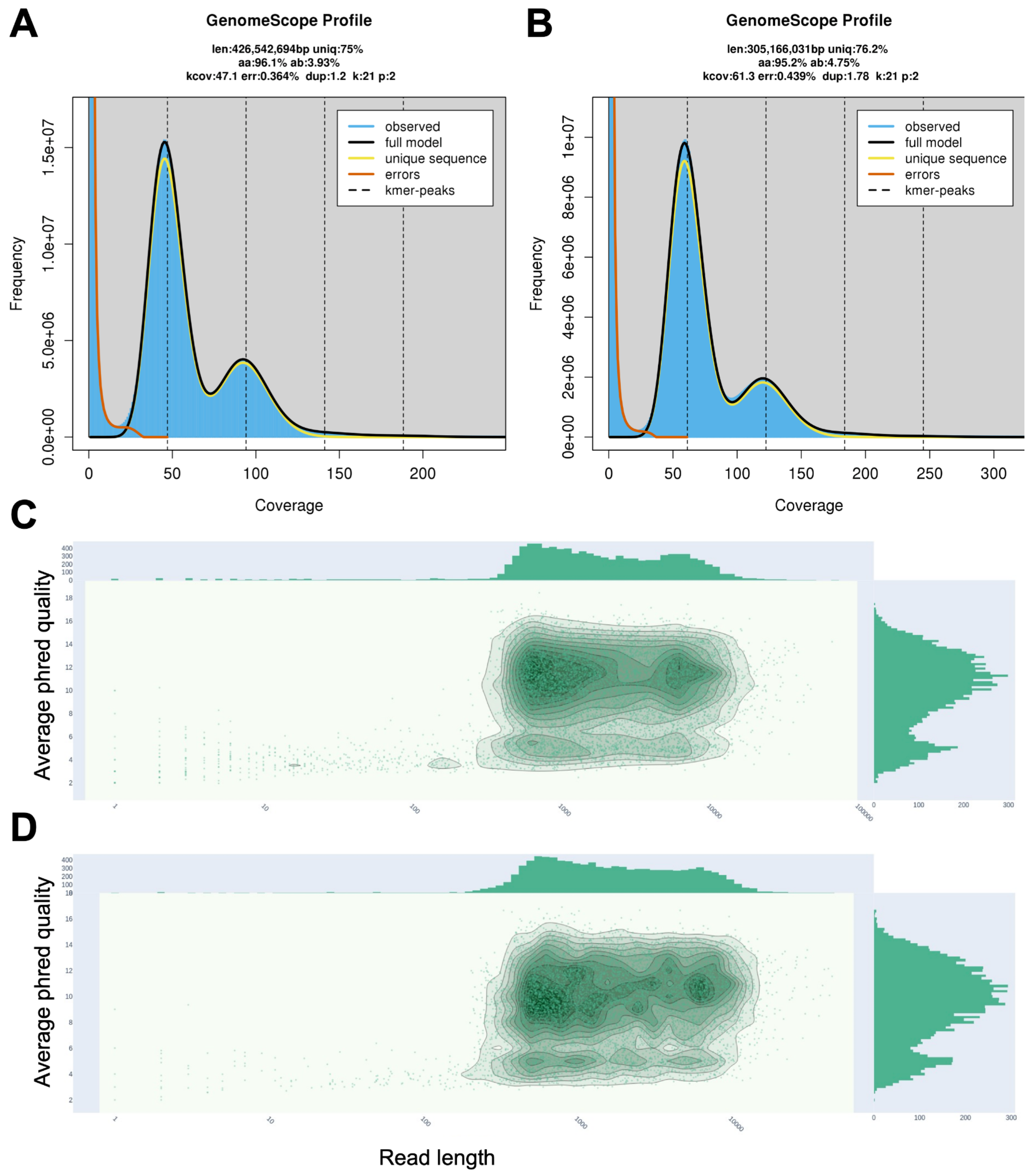


**Fig S1**. Statistics for Illumina (short reads) and ONT sequencing. GenomeScope analysis of Illumina sequencing reads of (A) *Asymmetron* and (B) *Epigonichthys*. Bivariant plot using NanoPlot comparing log-transformed ONT read length with average basecall Phred quality score for (A) *Asymmetron* and (B) *Epigonichthys*.

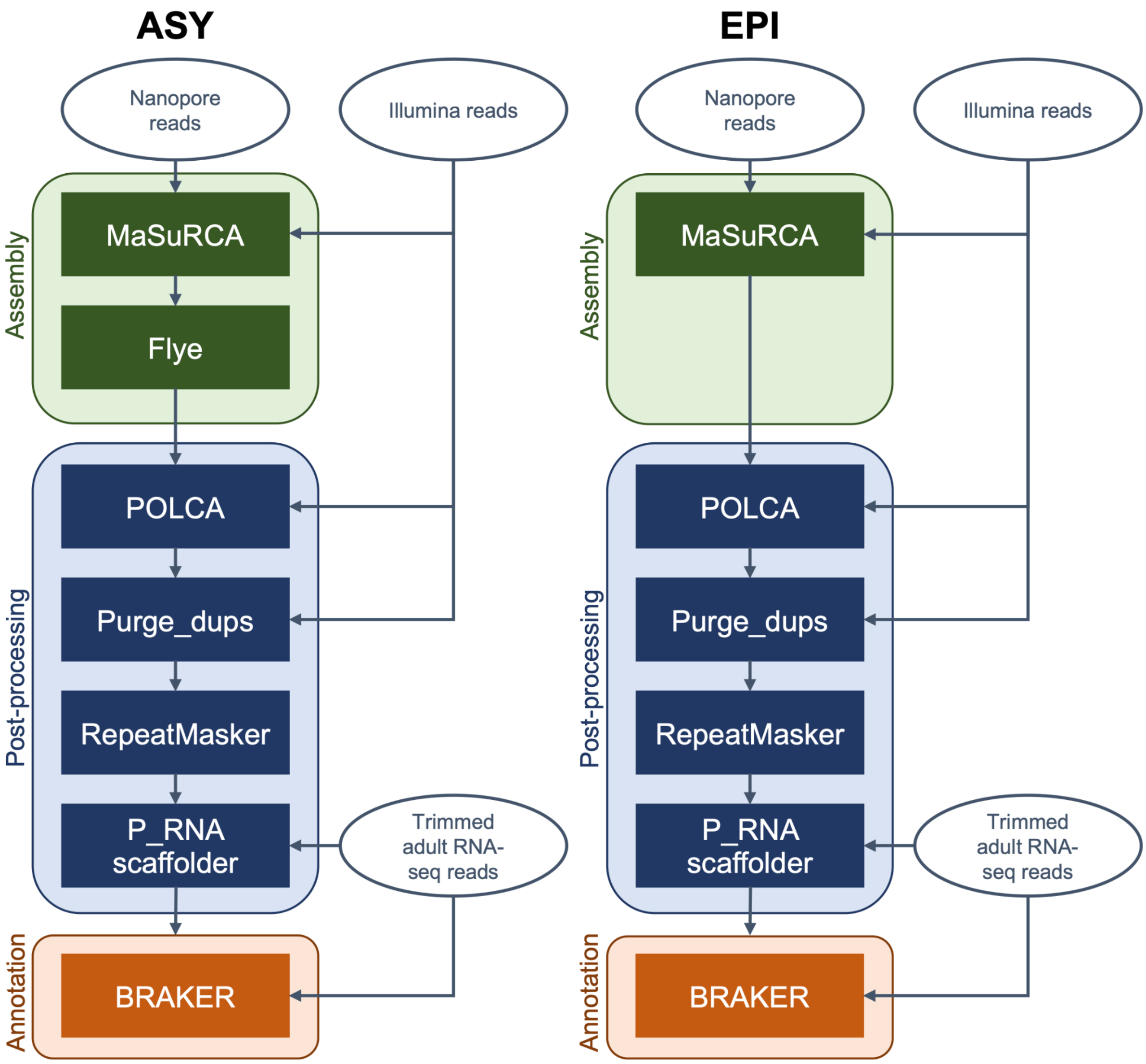


**Fig S2**. Assembly approaches and tools used to generate the final draft genomes and annotations of *Asymmetron* and *Epigonichthys*.

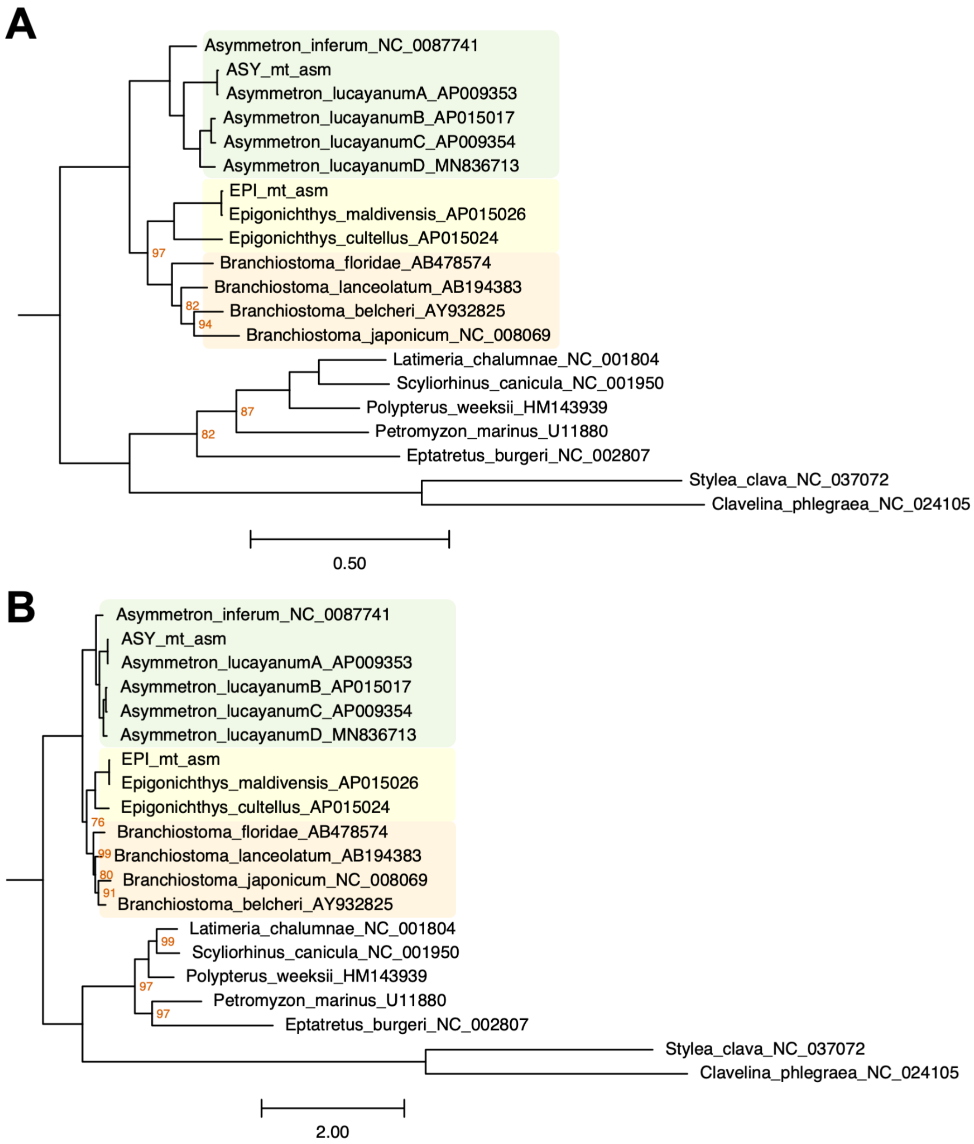


**Fig S3**. Mitogenome phylogenies based on (A) partitioned analysis using a best-partitioning scheme found within IQ-tree and (B) C60 mixture model within IQ-tree support the Asymmetron-sister topology. Nodes without support labels have maximal support.

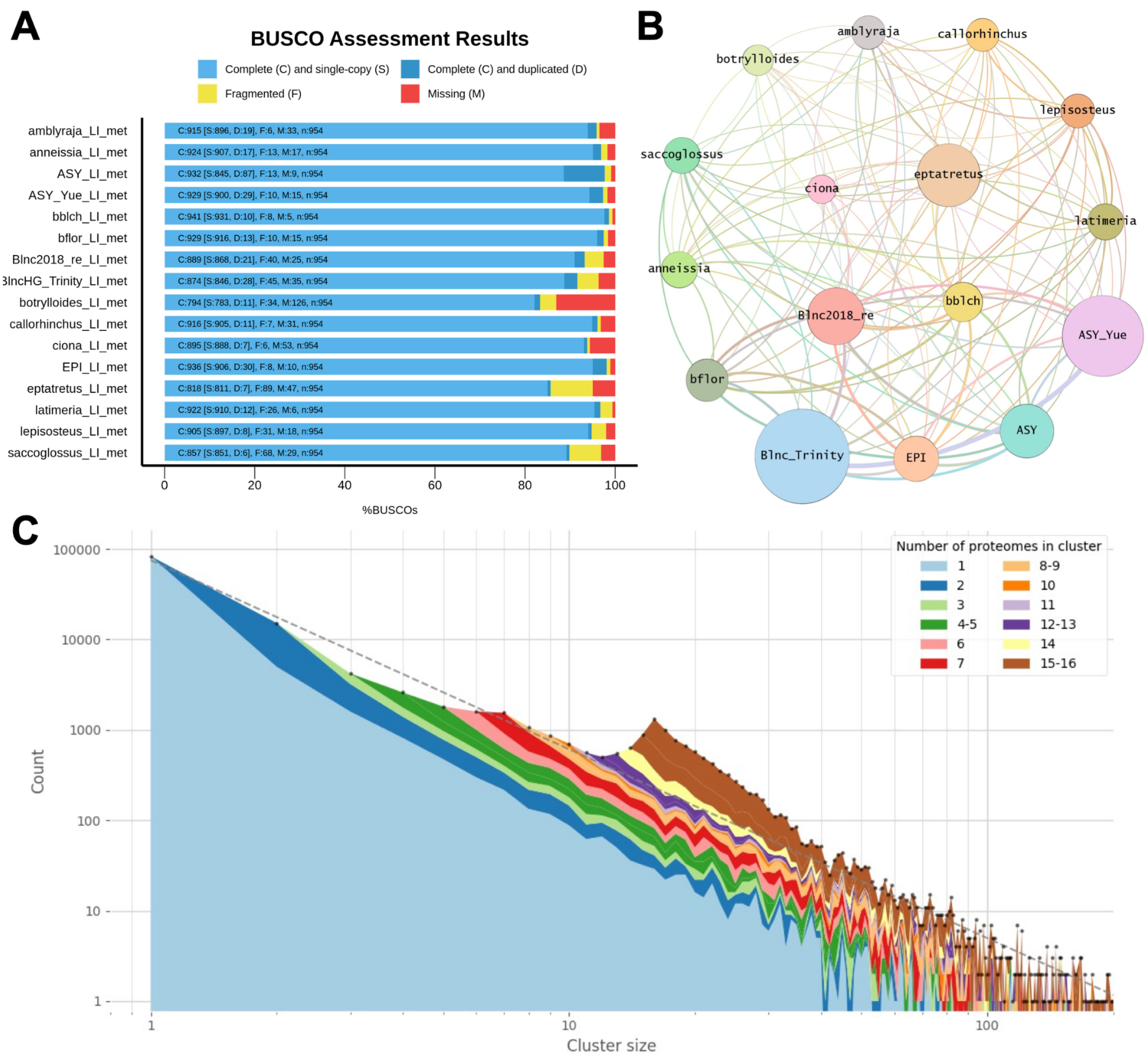


**Fig S4**. The proteome dataset used in the phylogenetic analysis. (A) High overall BUSCO completeness and low duplicate score of the proteomes. (B) Network representation of the clustering. The proteome size of each taxon corresponds to the node radius. The edges are weighted based on the number of times two taxa share an orthogroup. (C) Distribution of orthogroup sizes filled with the number of taxa included.

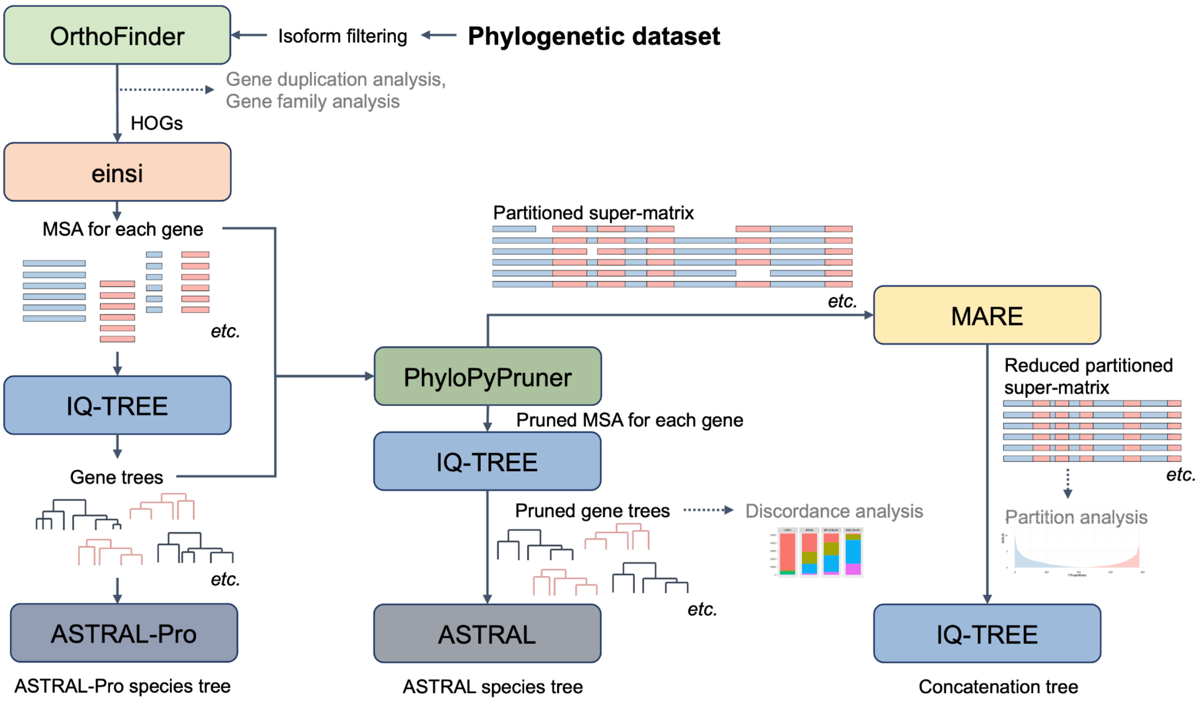


**Fig S5**. Phylogenetic inference approaches and tools used to generate the quartet-based ASTRAL and the concatenation-based IQ-TREE analyses. The associated analysis steps are also included.

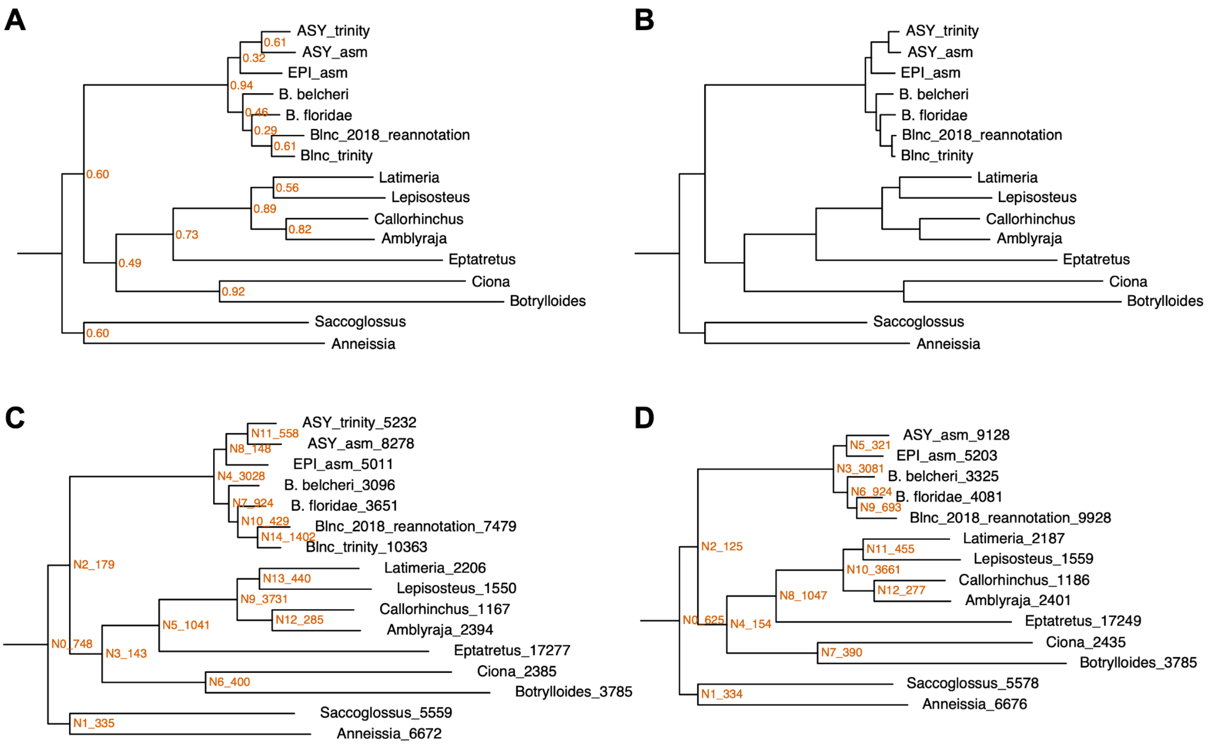


**Fig S6**. Default phylogenetic trees from OrthoFinder. (A) The automatically generated STAG tree supports the revised topology. The support values are based on the proportion of gene tree bipartitions that contain the dominant bipartition. (B) MSA tree (automatically generated using the option `-M msa`) supports the revised topology. All nodes have 100% bootstrap support. (C) Gene duplications as inferred from OrthoFinder are shown in the internal and terminal nodes. (D) The level of gene duplications at the base of cephalochordates remains stable after removing transcriptome assemblies (that may inflate the number of genes).

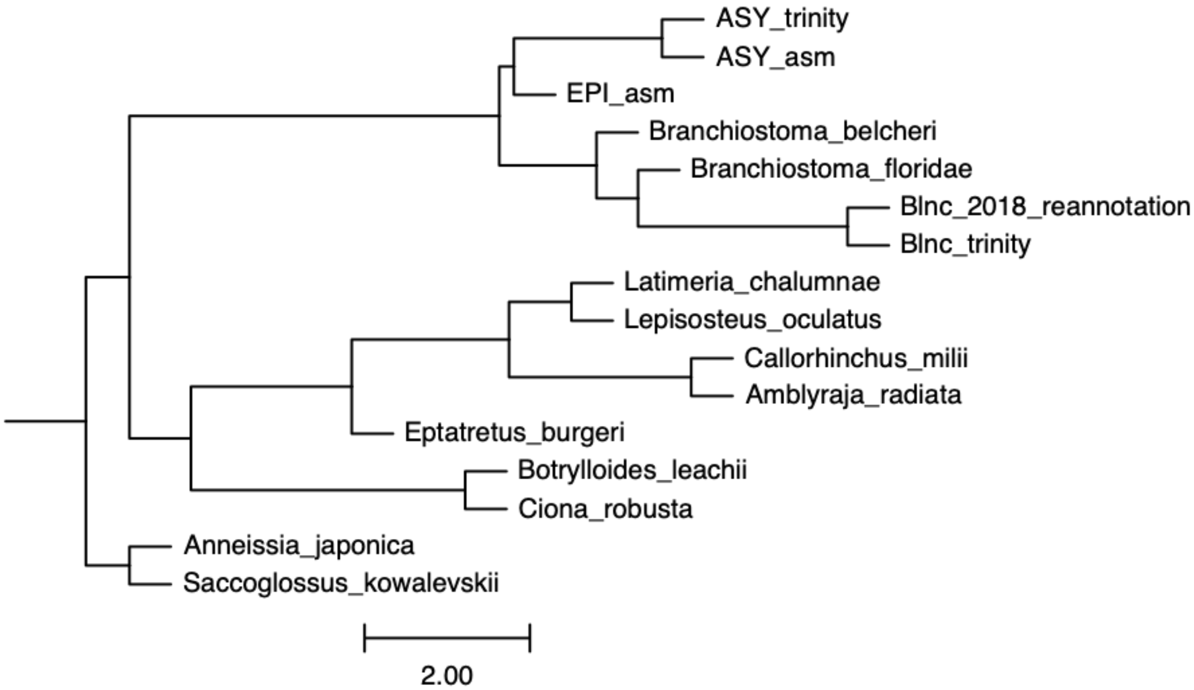


**Fig S7**. Paralogue-aware ASTRAL-Pro places *Branchiostoma* as a sister group to *Asymmetron* and *Epigonichthys*. All branches have maximal support.

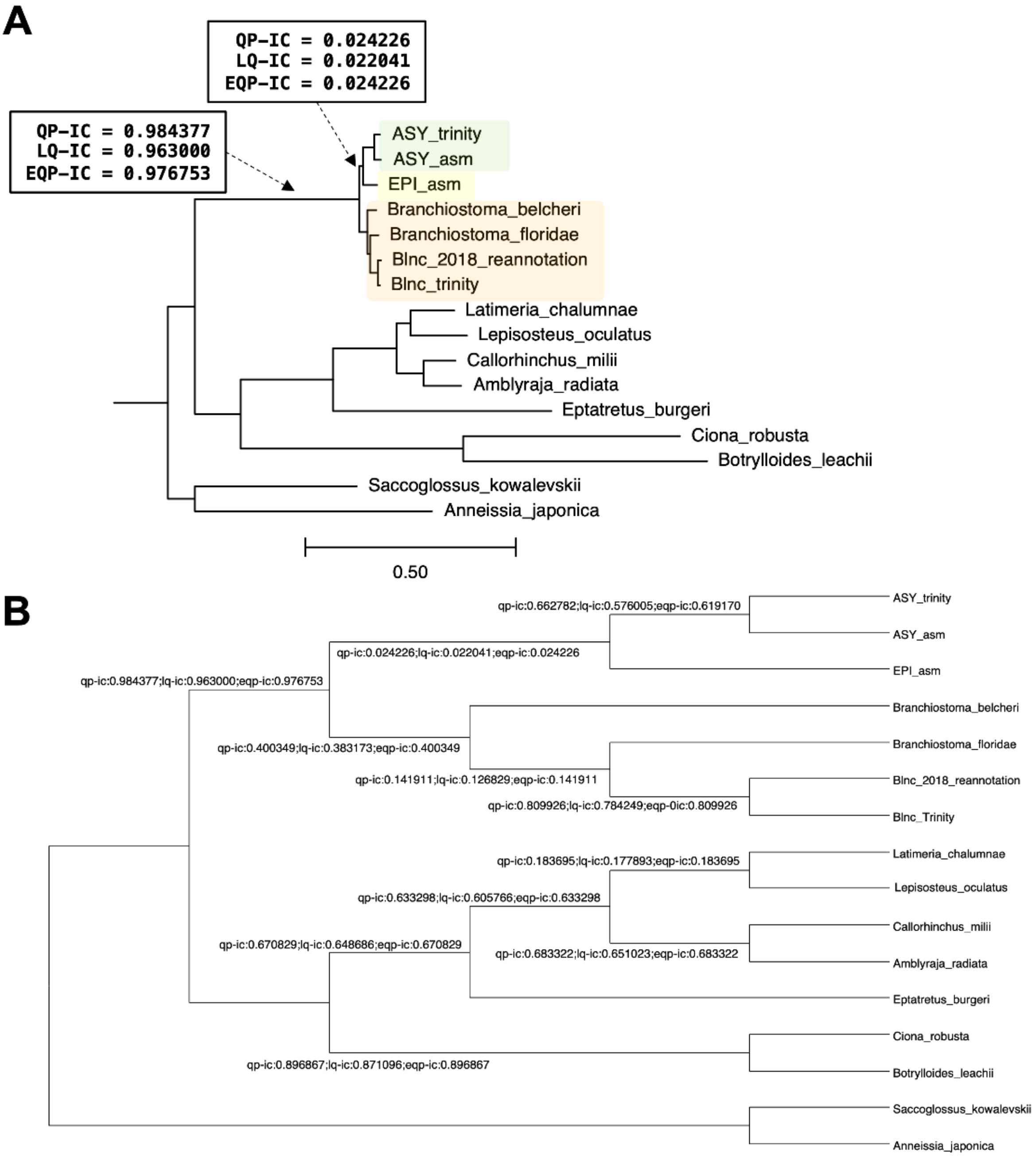


**Fig S8**. Internode certainty scores (A) at the branch leading to cephalochordates and the *Asymmetron*-*Epigonichthys* sistergroup, and (B) across the phylogenetic dataset. Dendroscope (Huson *et al.*, 2007) was used to visualise the internode certainty scores in (B).

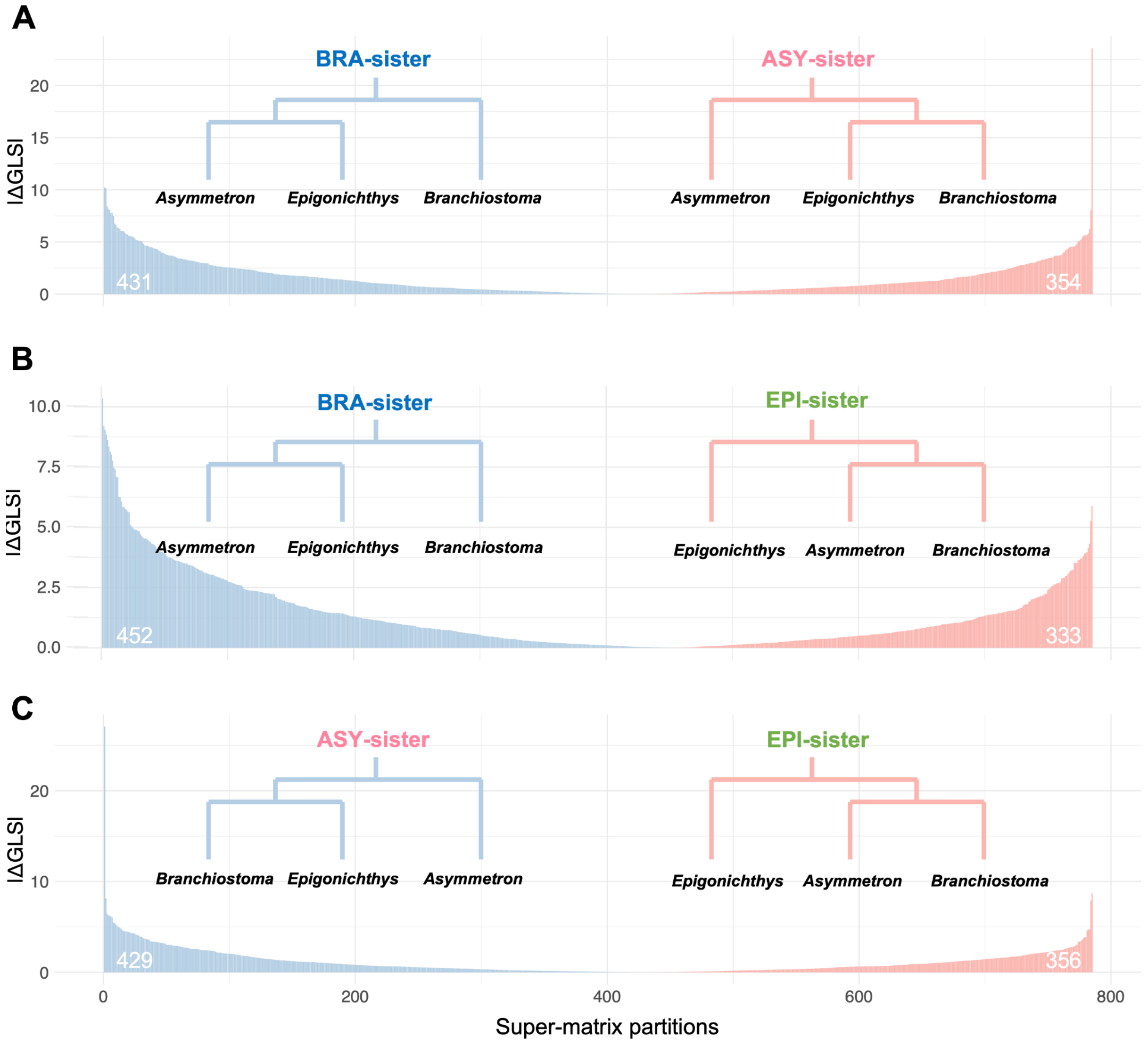


**Fig S9**. Gene-wise log-likelihood signal for each topology at each partition. The numbers on the plot indicate the number of partitions that support each topology.

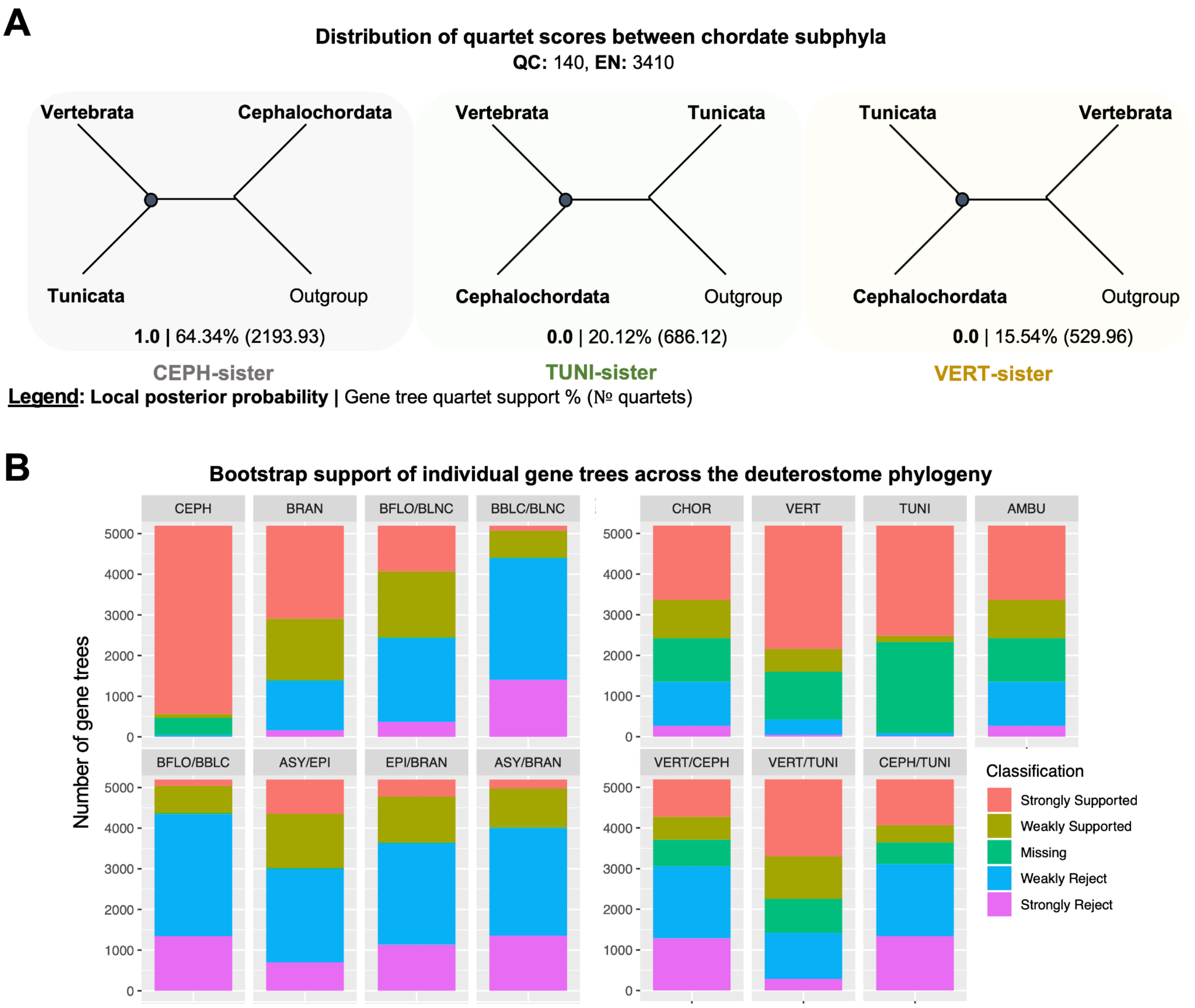


**Fig S10**. Gene tree discordance in other branches of the species tree. (A) The distribution of quartet scores used in the ASTRAL analysis between the three possible topologies for chordate subphyla. Cephalochordate-sister (Aa) has full local posterior probability support and is supported by a majority of gene tree quartets, although a large proportion of quartet also support tunicate-sister (Ab) and vertebrate-sister (Ac). (B) Analysis of the bootstrap support of the individual gene trees that support each grouping further demonstrates the existence of gene tree discordance across the deuterostome tree of life, although this effect is most prevalent amongst amphioxus genera.

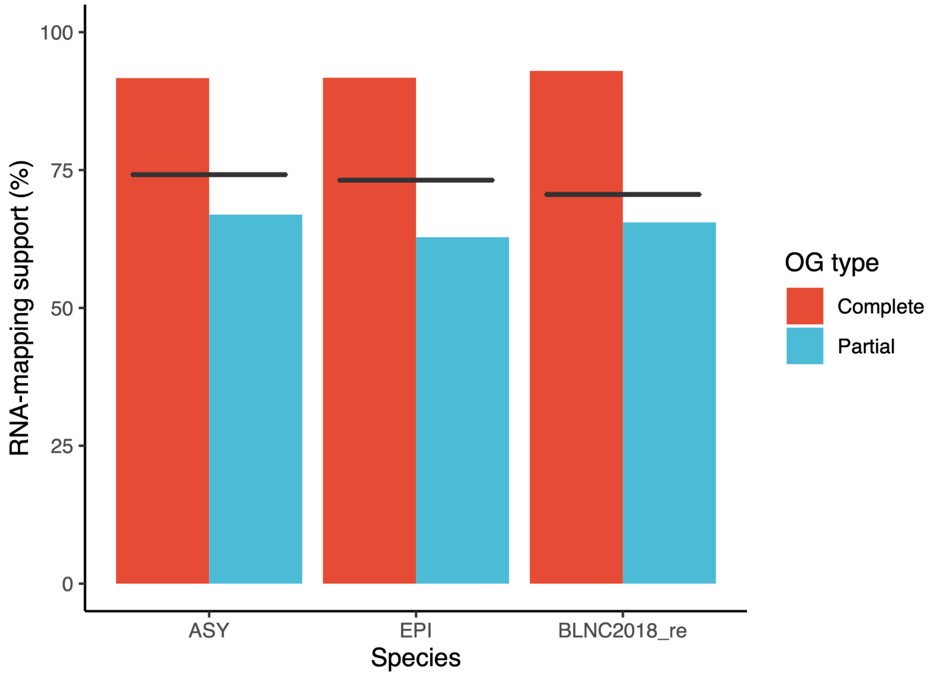


**Fig S11**. Percentage of gene models with RNA-seq mapping in cephalochordate-exclusive orthogroups (OG) from *Asymmetron* and *Epigonichthys* gene annotations and the *B. lanceolatum* gene re-annotation. Black lines denote the RNA-mapping support for all gene models in the respective annotations (regardless of OG membership).

**A**

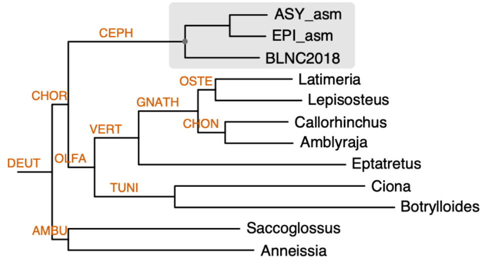


| Node name | Gene duplications | Exclusive orthogroups (complete / partial / all) | Gene family evolution (expansion / contraction) | Proteome count |
|---|---|---|---|---|
| **DEUT** | 643 | 4382 / 6194 / 10576 | NA | 12 |
| **AMBU** | 324 | 301 / 0 / 301 | +97 / -11 | 2 |
| **CHOR** | 170 | 53 / 939 / 992 | +0 / -33 | 10 |
| **OLFA** | 125 | 41 / 160 / 201 | +9 / -1265 | 7 |
| **CEPH** | 3254 | 3609 / 1224 / 4833 | +1172 / -1038 | 3 |
| **VERT** | 1006 | 376 / 241 / 617 | +1065 / -633 | 5 |
| **TUNI** | 405 | 669 / 0 / 669 | +84 / -2181 | 2 |
| **GNATH** | 3663 | 595 / 444 / 1039 | +1529 / -496 | 4 |
| **CHON** | 282 | 174 / 0 / 174 | +151 / -507 | 2 |
| **OSTE** | 422 | 144/ 0 / 144 | +233 / -132 | 2 |

**B**

**3 discrete gamma rate categories**

| Node name | Gene family evolution (expansion / contraction) |
|---|---|
| **DEUT** | NA |
| **AMBU** | +100 / -17 |
| **CHOR** | +1 / -29 |
| **OLFA** | +9 / -1268 |
| **CEPH** | +1184 / -1024 |
| **VERT** | +1067 / -629 |
| **TUNI** | +94 / -2180 |
| **GNATH** | +1532 / -495 |
| **CHON** | +153 / -501 |
| **OSTE** | +243 / -134 |

**C**

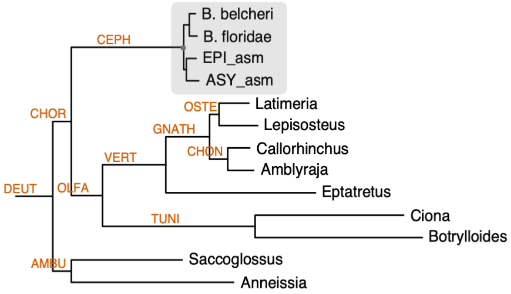


| Node name | Gene family evolution (expansion / contraction) |
|---|---|
| **DEUT** | NA |
| **AMBU** | +85 / -17 |
| **CHOR** | +2 / -60 |
| **OLFA** | +8 / -1253 |
| **CEPH** | +1263 / -1303 |
| **VERT** | +1113 / -681 |
| **TUNI** | +166 / -2362 |
| **GNATH** | +1505 / -467 |
| **CHON** | +153 / -605 |
| **OSTE** | +252 / -143 |

**Fig S12**. The level of genic innovations at the base of cephalochordates remain stable after removing datasets that may contain an inflated number of genes. (A) Calculated at the given nodes are: the gene duplications with at least 50% retention of duplicates in daughter lineages; orthogroups that are exclusive to the given node with 100% presence in daughter lineages (complete), with <100% presence (partial) and ≤100% presence (all); gene family expansion and contraction at the given node as estimated using cafe5 (lambda with no among family rate variation); and the number of proteomes at the given node. (B) Gene family expansion and contraction at the given node from (A) as estimated using cafe5 with 3 discrete gamma rate categories. (C) Gene family expansion and contraction at the given node as estimated using cafe5 (lambda with no among family rate variation), using another set of *Branchiostoma* datasets. For the cafe5 analysis, no orthogroups were removed.

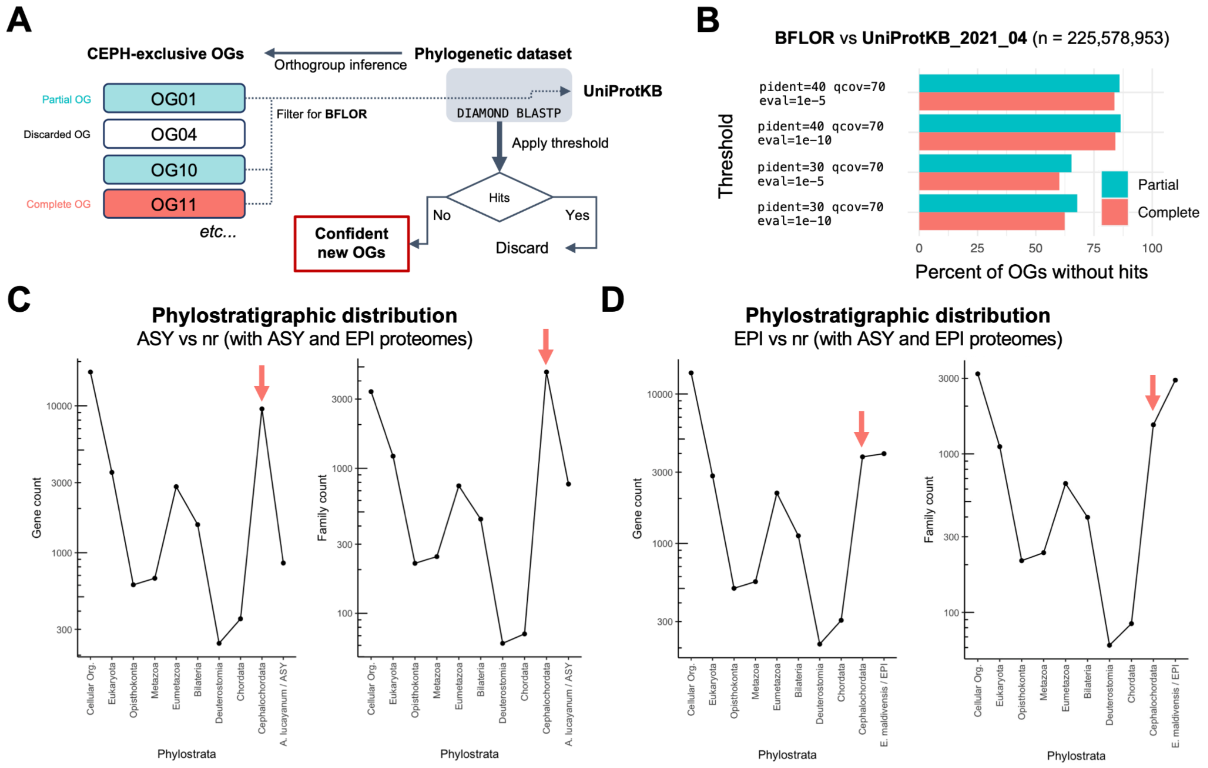


**Fig S13**. Further evidence of genic innovations. (A) Summary of the method used to assess whether any genes in the cephalochordate-exclusive orthogroups (i.e. orthogroups inferred at the last common cephalochordate ancestor) have any sequence homology outside of the group. (B) The proportion of confident new OGs at different thresholds. Phylostratigraphic distribution at the individual gene and inferred gene family levels in (C) *Asymmetron* and (D) *Epigonichthys*. The cephalochordate-level genes represent ~25% of the *Asymmetron* genome and ~13% of the *Epigonichthys* genome. The cephalochordate-level families represent ~39% of *Asymmetron* and ~15% of *Epigonichthys* gene families.

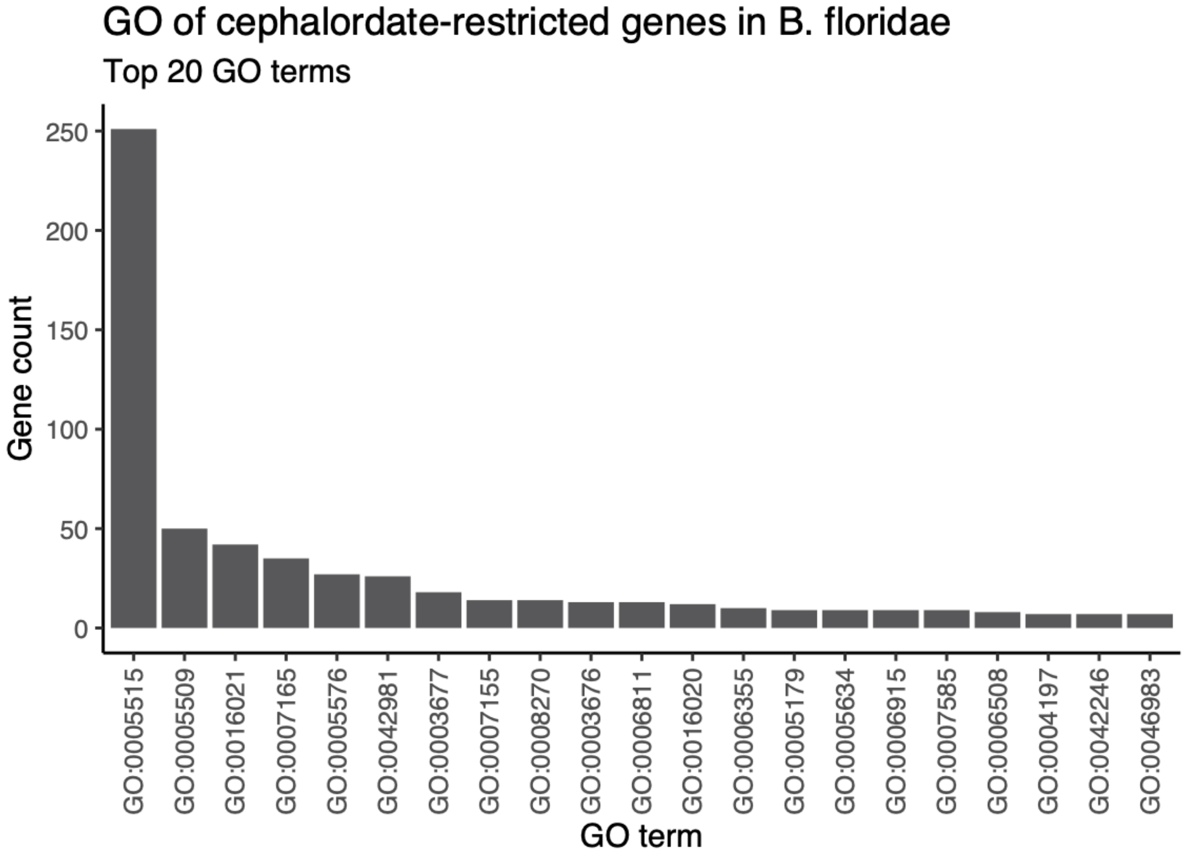


**Fig S14**. Most common gene ontology (GO) terms of *B. floridae* genes restricted to the cephalochordate phylostratum. For visualisation purposes, the top 20 GO terms are shown.

### Supplementary tables

**Table S1**. Tested tools and parameters for assembling *Asymmetron* and *Epigonichthys* genomes. Unless otherwise stated, default parameters were used for each assembly. For Canu, “keep haplotypes” and “collapse haplotypes” correspond to the parameters `corOutCoverage=200 "batOptions=-dg 3 -db 3 -dr 1 -ca 500 -cp 50"` and `corOutCoverage=200 correctedErrorRate=0.15`, respectively. The parameters from MaSuRCA's customised Flye v2.5 which were used in Flye v2.8.2 are explained in the method section. BUSCO's eukaryotic database from OrthoDB v10, consisting of 255 single-copy orthologues, was used to assess completeness rather than the metazoan database due to faster run-time. The assemblies ultimately used after post-processing are emboldened.

| Tools (options) | Assembly statistics | | | |
|---|---|---|---|---|
| | Length (Mb) | № contigs | N50 (kb) | BUSCO |
| *Asymmetron* (putative *A. lucayanum* clade A) | | | | |
| Canu | 656 | 20,843 | 47.0 | C:73.7% [S:63.1%] |
| Canu (keep haplotypes) | 705 | 21,902 | 47.4 | C:71.4%[S:60.8%] |
| Flye | 628 | 33,568 | 60.7 | C:88.2%[S:80.4%] |
| Flye (`--keep-haplotypes`) | 592 | 46,299 | 36.8 | C:73.7%[S:69.0%] |
| Flye (`--meta`) | 628 | 29,364 | 68.2 | C:87%[S:78.4%] |
| Flye (`--meta` & `--keep-haplotypes`) | 598 | 29,364 | 43.9 | C:82.85[S:76.1%] |
| NextDenovo | 351 | 3,498 | 122.3 | C:60.8%[S:60.4%] |
| Haslr | 309 | 37,408 | 13.8 | C:22.4%[S:22.0%] |
| MaSuRCA | 793 | 13,755 | 85.5 | C:95.7%[S:38.4%] |
| MaSuRCA (`LHE_COVERAGE=60`) | 795 | 13,834 | 85.2 | C:96.1%[S:39.2%] |
| MaSuRCA (`LHE_COVERAGE=60`) & Flye | 836 | 43,847 | 50.5 | C:96.9%[S:40.8%] |
| MaSuRCA (`LHE_COVERAGE=60`) & Flye (`--keep-haplotypes`) | 761 | 64,271 | 32.2 | C:87.0%[S:34.5%] |
| **MaSuRCA (`LHE_COVERAGE=60`) & Flye (MaSuRCA parameters)** | **854** | **28,435** | **77.4** | **C:95.3%[S:34.9%]** |
| MaSuRCA & Canu | 803 | 18,493 | 71.1 | C:96.1%[S:38.8%] |
| MaSuRCA (`LHE_COVERAGE=60`) & Canu | 802 | 18,482 | 71.3 | C:96.0%[S:38.4%] |

| | | | | |
|---|---|---|---|---|
| Platanus-Allee | 650 | 536,791 | 60.0 | C:93.7%[S:83.9%] |
| *Epigonichthys* (putative *E. maldivensis*) | | | | |
| Canu | 582 | 12,958 | 78.2 | C:88.7%[S:66.3%] |
| Canu (keep haplotypes) | 624 | 9,969 | 117.8 | C:89.4%[S:58.8%] |
| Canu (collapse haplotypes) | 572 | 11,723 | 87.7 | C:87.9%[S:65.9%] |
| Flye | 458 | 26,420 | 72.8 | C:95.3%[S:85.5% |
| Flye (`--keep-haplotypes`) | 421 | 34,950 | 38.9 | C:80.8%[S:71.8%] |
| Flye (`--meta`) | 455 | 22,263 | 80.2 | C:91.0%[S:74.9%] |
| Flye (`--meta` & `--keep-haplotypes`) | 431 | 25,981 | 51.2 | C:89.8%[S:74.5%] |
| NextDenovo | 329 | 1,125 | 832.8 | C:88.6%[S:88.6%] |
| Haslr | 268 | 21,249 | 22.3 | C:45.1%[S:38.8%] |
| MaSuRCA | 613 | 5,803 | 230.4 | C:98.9%[S:22.4%] |
| **MaSuRCA (`LHE_COVERAGE=60`)** | **608** | **6,118** | **209.9** | **C:98.8%[S:20.8%]** |
| MaSuRCA & Flye | 636 | 29,058 | 84.7 | C:95.6%[S:27.8%] |
| MaSuRCA & Flye (`--keep-haplotypes`) | 566 | 39,780 | 57.4 | C:89.4%[S:20.0%] |
| MaSuRCA (`LHE_COVERAGE=60`) & Flye | 636 | 28,671 | 86.0 | C:95.7%[S:26.7%] |
| MaSuRCA (`LHE_COVERAGE=60`) & Flye (`--keep-haplotypes`) | 567 | 39,509 | 58.3 | C:90.6%[S:21.2%] |
| MaSuRCA & Flye (MaSuRCA parameters) [EPI_3] | 640 | 14,731 | 170.3 | C:97.7%[S:21.2%] |
| MaSuRCA (`LHE_COVERAGE=60`) & Flye (MaSuRCA parameters) | 640 | 14,807 | 164.0 | C:98.5%[S:21.6%] |
| MaSuRCA & Canu | 616 | 9,146 | 143.1 | C:98.8%[S:23.5%] |
| MaSuRCA (`LHE_COVERAGE=60`) & Canu | 615 | 9,122 | 144.0 | C:99.2%[S:23.1%] |
| Platanus-Allee | 441 | 207,096 | 117.6 | C:97.2%[S:83.1%] |

Tools used:

- Canu v2.1 (Koren *et al.*, 2017)
- Flye v2.8.2 (Kolmogorov *et al.*, 2019)
- NextDenovo v2.4.0 (https://github.com/Nextomics/NextDenovo)
- MaSuRCA v3.4.2 (Zimin *et al.*, 2017)
- HASLR v0.8a1 (Haghshenas *et al.*, 2020)
- Platanus-Allee v2.2.2 (Kajitani *et al.*, 2019)

**Table S2**. Post-processing improves the *Asymmetron* and *Epigonichthys* assemblies. The BUSCO completeness was analysed with both eukaryota (255 single-copy orthologues) and metazoan datasets (954 single-copy orthologues) from OrthoDB v10. The metazoan results are italicised. It should be noted that the assemblies chosen here were compared with the post-processing of other outputs from the assembly step before being chosen.

| Tools (options) | Assembly statistics | | | |
|---|---|---|---|---|
| | Length (Mb) | № contigs | N50 (kb) | BUSCO |
| *Asymmetron* (putative *A. lucayanum* clade A) | | | | |
| Polishing: POLCA | 855 | 28,435 | 77.4 | C:97.7%[S:31.0%] |
| Haplotig-purging: purge_dups (short-reads, manual cutoffs, asm20, no -e) | 512 | 8,942 | 115.5 | C:96.5%[S:90.2%] |
| Repeatmasking + scaffolding: RepeatMasker/RepeatModeler | 512 | 6,488 | 207.6 | C:97.7%[S:92.2%] *C:97.0%[S:90.9%]* |
| *Epigonichthys* (putative *E. maldivensis*) | | | | |
| Polishing: POLCA | 608 | 6,118 | 210.0 | C:98.9%[S:17.3%] |
| Haplotig-purging: purge_dups (short-reads, manual cutoffs, asm20, no -e) | 345 | 2,595 | 325.1 | C:98.5%[S:96.5%] |
| Repeatmasking + scaffolding: RepeatMasker/RepeatModeler | 345 | 1,808 | 564.7 | C:99.3%[S:97.3%] *C:97.6%[S:95.3%]* |

Tools used:

- POLCA (Zimin and Salzberg, 2020)
- Purge_dups v1.2.5 (Guan *et al.*, 2020)
- RepeatModeler v2.0.1 (Flynn *et al.*, 2020)
- RepeatMasker v4.1.2 (https://www.repeatmasker.org/)
- P_RNA_scaffolder (Zhu *et al.*, 2018)

**Table S3.** Summary of the genome assemblies, transcriptome assemblies and re-annotation used in the phylogenetic dataset. All taxa here belong to subphylum Cephalochordata.

| Steps | Tools, parameters and other notes |
|---|---|
| **ASY_asm** - *Asymmetron* (putative *A. lucayanum* clade A) | |
| Assembly | MaSuRCA (option:`LHE_COVERAGE=60`) & Flye (MaSuRCA parameters) |
| Polishing | POLCA (short reads) |
| Haplotig-purging | purge_dups (short reads pipeline, `-xasm 20`, manual cutoffs, and no `-e`) |
| Repeat-masking | RepeatModeler/RepeatMasker |
| RNA-scaffolding | P_RNA_scaffolder (see method section for detail) |
| Annotation | BRAKER ET mode |
| **EPI_asm** - *Epigonichthys* (putative *E. maldivensis*) | |
| Assembly | MaSuRCA (option:`LHE_COVERAGE=60`) |
| Polishing | POLCA (short reads) |
| Haplotig-purging | purge_dups (short reads pipeline, `-xasm 20`, manual cutoffs, and no `-e`) |
| Repeat-masking | RepeatModeler/RepeatMasker |
| RNA-scaffolding | P_RNA_scaffolder (see method section for detail) |
| Annotation | BRAKER ET mode |
| **ASY_trinity** - *A. lucayanum* (from Yue et al. 2014) | |
| Transcriptome re-assembly | NCBI SRA accession: SRX437621 (Adult)<br>Trinity (genome-free mode) |
| **Blnc_trinity** - *B. lanceolatum* (another Mediterranean isolate) | |
| Transcriptome assembly | Trinity (genome-guided mode) |
| **Blnc_2018_reannotation** - *B. lanceolatum* (from Marlétaz et al. 2018) | |
| Re-annotation | GenBank assembly accession GCA_900088365<br>BRAKER ET mode |

**Table S4**. Species in the phylogenetic dataset and source.

| Species | Source |
| --- | --- |
| *Asymmetron* (putative *A. lucayanum* clade A) [ASY_asm] | Genome assembly and annotation from this study |
| *Epigonichthys* (putative *E. maldivensis*) [EPI_asm] | Genome assembly and annotation from this study |
| *A. lucayanum* (Trinity) [ASY_trinity] | Transcriptome assembly of NCBI SRA: SRX437621 |
| *Branchiostoma lanceolatum* (Trinity) [Blnc_trinity] | Transcriptome assembly from this study |
| *Branchiostoma lanceolatum* (re-annotation) [Blnc_2018_reannotation] | Re-annotation of GenBank assembly accession: GCA_900088365 |
| *Amblyraja radiata* | RefSeq: GCF_010909765.2 |
| *Anneissia japonica* | RefSeq: GCF_011630105.1 |
| *Branchiostoma belcheri* | RefSeq: GCF_001625305.1 |
| *Branchiostoma floridae* | RefSeq: GCF_000003815.2 |
| *Botrylloides leachii* | www.aniseed.cnrs.fr/aniseed/ |
| *Callorhinchus milii* | RefSeq: GCF_000165045.1 |
| *Ciona robusta* | RefSeq: GCF_000224145.3 |
| *Eptatretus burgeri* | www.figshare.com/projects/eburgeri-genome/77052 |
| *Latimeria chalumnae* | RefSeq: GCF_000225785.1 |
| *Lepisosteus oculatus* | RefSeq: GCF_000242695.1 |
| *Saccoglossus kowalevskii* | RefSeq: GCF_000003605.2 |

**Table S5**. Transcriptome assembly of the Mediterranean isolate of *B. lanceolatum* and the publicly available *A. lucayanum* RNA-seq dataset. `Genome-guided' and `genome-free' refer to Trinity assembly with and without a genome that is used to guide the transcriptome assembly. The BUSCO completeness was analysed using the metazoan datasets (954 single-copy orthologues) from OrthoDB v10. The assemblies ultimately used are emboldened.

| **Trinity options** | **Assembly statistics** | | | |
|---|---|---|---|---|
| | **№ transcripts** | **N50 (bp)** | **Ex90N50 (bp)** | **BUSCO** |
| *B. lanceolatum* (Mediterranean isolate) | | | | |
| Genome-free | 594,688 | 1504 | 1443 | C:98.6%[S:6.9%] |
| **Genome-guided** | **332,492** | **2084** | **1924** | **C:98.1%[S:9.4%]** |
| *A. lucayanum* (SRX437621) | | | | |
| **Genome-free (adult RNA-seq)** | **191,443** | **2038** | **2074** | **C:98.6%[S:11.8%]** |
| Genome-free (adult + embryo RNA-seq) | 565,148 | 1291 | 1412 | C:98.8%[S:8.7%] |

**Table S6**. Isoform filtering reduces the proteome size and the BUSCO duplicate scores while maintaining the overall completeness for most taxa. The BUSCO completeness was analysed using the metazoan datasets (954 single-copy orthologues) from OrthoDB v10.

| Species | Assembly statistics | | | |
|---|---|---|---|---|
| | № protein sequences | | BUSCO | |
| | Pre-filtered | Filtered | Pre-filtered | Filtered |
| *Asymmetron* (putative *A. lucayanum* clade A) [ASY_asm] | 45,562 | 37,374 | C:97.7%[S:73.0%] | C:97.7%[S:88.6%] |
| *Epigonichthys* (putative *E. maldivensis*) [EPI_asm] | 35,357 | 29,547 | C:98.1%[S:82.1%] | C:98.1%[S:95.0%] |
| *A. lucayanum* (Trinity) [ASY_trinity] | 179,017 | 63,373 | C:98.1%[S:12.4%] | C:97.3%[S:94.3%] |
| *Branchiostoma lanceolatum* (Trinity) [Blnc_trinity] | 247,134 | 76,186 | C:97.8%[S:10.9%] | C:91.6%[S:88.7%] |
| *Branchiostoma lanceolatum* (re-annotation) [Blnc_2018_reannotation] | 51,037 | 40,855 | C:93.2%[S:73.4%] | C:93.2%[S:91.0%] |
| *Amblyraja radiata* | 38,895 | 18,474 | C:96.1%[S:57.9%] | C:95.9%[S:93.9%] |
| *Anneissia japonica* | 32,774 | 21,084 | C:97.0%[S:70.0%] | C:96.9%[S:95.1%] |
| *Branchiostoma belcheri* | 34,675 | 23,855 | C:98.7%[S:75.3%] | C:98.6%[S:97.6%] |
| *Branchiostoma floridae* | 43,041 | 26,676 | C:97.4%[S:72.6%] | C:97.4%[S:96.0%] |
| *Botrylloides leachii* | 15,839 | 15,833 | C:83.3%[S:82.1%] | C:83.3%[S:82.1%] |
| *Callorhinchus milii* | 28,237 | 17,843 | C:96.1%[S:75.8%] | C:96.1%[S:94.9%] |
| *Ciona robusta* | 21,096 | 13,713 | C:94.2%[S:79.9%] | C:93.8%[S:93.1%] |
| *Eptatretus burgeri* | 50,127 | 46,261 | C:86.1%[S:71.0%] | C:85.7%[S:85.0%] |
| *Latimeria chalumnae* | 34,251 | 20,932 | C:96.9%[S:68.4%] | C:96.7%[S:95.4%] |
| *Lepisosteus oculatus* | 41,647 | 18,771 | C:94.9%[S:65.3%] | C:94.8%[S:94.0%] |
| *Saccoglossus kowalevskii* | 22,111 | 20,922 | C:90.0%[S:84.4%] | C:89.8%[S:89.2%] |

**Table S7**. Summary of the orthology assignment.

| Statistic | Count |
|---|---|
| Number of species | 16 |
| Number of genes | 491699 |
| Number of genes in orthogroups | 408482 |
| Number of unassigned genes | 83217 |
| Percentage of genes in orthogroups | 83.1 |
| Percentage of unassigned genes | 16.9 |
| Number of orthogroups | 41723 |
| Number of species-specific orthogroups | 9357 |
| Number of genes in species-specific orthogroups | 42901 |
| Percentage of genes in species-specific orthogroups | 8.7 |
| Mean orthogroup size | 9.8 |
| Median orthogroup size | 4.0 |
| G50 (assigned genes) | 19 |
| G50 (all genes) | 16 |
| O50 (assigned genes) | 5669 |
| O50 (all genes) | 8068 |
| Number of orthogroups with all species present | 4255 |
| Number of single-copy orthogroups | 658 |
